# Direct observation of fluorescent proteins in gels: a rapid cost-efficient, and quantitative alternative to immunoblotting

**DOI:** 10.1101/2024.05.31.594679

**Authors:** Matthieu Sanial, Ryan Miled, Marine Alves, Sandra Claret, Nicolas Joly, Véronique Proux-Gillardeaux, Anne Plessis, Sébastien Léon

## Abstract

The discovery of Green Fluorescent Protein (GFP) and its derivatives has revolutionized cell biology. These fluorescent proteins (FPs) have enabled the real-time observation of protein localization and dynamics within live cells. Applications of FP vary from monitoring gene/protein expression patterns, visualizing protein-protein interactions, measuring protein stability, assessing protein mobility and creating biosensors. The utility of FPs also extends to biochemical approaches through immunoblotting and proteomic analyses, aided by anti-FP antibodies and nanobodies. FPs are notoriously robust proteins with a tightly folded domain that confers a strong stability and a relative resistance to degradation and denaturation. In this study, we report that various green, orange and red FPs can be maintained in a native, fluorescent state during the entire process of protein sample extraction, incubation with sample buffer, loading and migration on SDS-PAGE with only minor adaptations of traditional protocols. This protocol results in the ability to detect and quantify in-gel fluorescence (IGF) of endogenously-expressed proteins tagged with FPs directly after migration, using standard fluorescence-imaging devices. This approach eliminates the need for antibodies and chemiluminescent reagents, as well as the time-consuming steps inherent in immunoblotting such as transfer onto a membrane and antibody incubations. Overall, IGF detection provides clearer data with less background interference, a sensitivity comparable or better to antibody-based detection, a better quantification and a broader dynamic range. After fluorescence imaging, gels can still be used for other applications such as total protein staining or immunoblotting if needed. It also expands possibilities by allowing the detection of FPs for which antibodies are not available. Our study explores the feasibility, limitations, and applications of IGF for detecting endogenously expressed proteins in cell extracts, providing insights into sample preparation, imaging conditions, and sensitivity optimizations, and potential applications such as co-immunoprecipitation experiments.

## Introduction

The development of GFP as a tool to study protein localization and dynamics in live cells has been a revolution in the field of cell biology, as recognized by the Nobel prize in Chemistry awarded in 2008 to Osamu Shimomura, Martin Chalfie and Roger Tsien (Chalfie et al., 1994; Tsien, 1998). GFP and its derivatives, as well as other fluorescent proteins (FPs) isolated since then from other organisms with various properties (Lambert, 2019; Shaner et al., 2005), have become instrumental in many fields of biology. They allow to monitor protein expression and localization in live cells, and can be used as reporters of gene/protein expression patterns in organisms or tissues. Various techniques have been implemented using GFP and its variants to visualize protein-protein interactions in cells, through bimolecular fluorescence complementation (Hu and Kerppola, 2003; Romei and Boxer, 2019), or fluorescence resonance energy transfer (FRET, BRET) (Hochreiter et al., 2015). This also opened the way to the development of various probes and biosensors to report on the quantity and/or localization of ions, metabolites and other organic molecules in cells and tissues (Chandris et al., 2021; Wang et al., 2023; Zacharias et al., 2002). Diverse folding kinetics of FPs have been exploited to report on protein stability and half-life (fluorescent timers) (Khmelinskii et al., 2012; Subach et al., 2009; Terskikh et al., 2000). The fact that fluorescence bleaches upon intense illumination also provides a way to evaluate protein mobility and dynamics in live cells (Fluorescence Recovery after Photobleaching [FRAP] and Fluorescence Loss in Photobleaching [FLIP]) (White and Stelzer, 1999). Similarly, photo-activatable and photo-switchable FPs allow the study of distinct protein pools over time by activating or converting their fluorescence upon illumination (Wang et al., 2023). Following the development of robust anti-GFP antibodies and in particular of nanobodies (Rothbauer et al., 2006), GFP can also be used like any other epitope for protein detection by techniques ranging from immunoblotting to immunoprecipitation and proteomic analyses (Cristea et al., 2005; Rothbauer et al., 2008). Finally, the ability to express a GFP-binding protein (derived from a llama single chain antibody) in cells fused with cellular proteins provides a way to modify the subcellular localization and interactions of a GFP-tagged protein at will (Rothbauer et al., 2008).

Because of these many applications, GFP and other FPs are frequently used as protein tags in cell biology studies. An early step in the workflow consists in verifying the expression of an intact fusion protein in cells by immunoblotting. Immunoblotting is also required when GFP is used as any epitope to monitor the fusion protein’s integrity, stability or post-translational modifications, or in interaction studies based on co-immunoprecipitations. Protein detection by immunoblotting involves primary antibodies and most generally a chemiluminescent reaction (ECL) using secondary antibodies coupled to an enzyme (horseradish peroxidase, HRP). Whereas this approach is considered to be more sensitive, HRP kinetics may interfere with the linearity of the signal vis-à-vis the amount of detected protein, a problem that may hinder the quantification of signals.

Over the past 20 years, fluorescent alternatives to chemiluminescence-based protein detection were introduced, and the development of fluorescent dyes and the amelioration of fluorescence-based imaging systems have allowed for a better sensitivity and better quantification of signals (Eaton et al., 2014). Although initial systems focused on fluorescence detection at high wavelengths (far-red and infrared, i.e. around 700 nm and above) to minimize background autofluorescence, commercial imaging systems now allow the visualization of fluorescence at additional wavelengths compatible with usual commercial fluorophores, eg. those used for immunofluorescence.

The versatility of these imaging systems is such that, in principle, the endogenous fluorescence of the widely used FPs (eg. GFP) in extracts could be monitored directly in gels without the need of transfer to membrane or the use of antibodies. SDS-PAGE involves denaturation of protein samples at high temperatures in the presence of SDS and reducing agents prior to loading onto a gel, procedures that are usually considered incompatible with the maintenance of FP fluorescence in gel (Chew et al., 2009). Other types of fluorescent proteins, such as using Flavin-binding FPs, bacteriophytochrome-based FPs, or bilirubin-binding FP (UnaG) can be used to detect proteins *in vivo* or *in vitro* (Rodriguez et al., 2017; Shcherbakova et al., 2015), some of which can be visualized in gel in the near-infrared from a fully-denatured extract in the presence of zinc acetate (Berkelman and Lagarias, 1986; Stepanenko et al., 2022). However, this relies on less commonly used fluorescent protein tags, with spectral properties that are not always available in classical fluorescence microscopy setup, usually preventing a combined use of these tags for imaging and biochemical studies.

A few studies have reported the observation of GFP-fluorescence in gels, but these approaches require the use of native PAGE (Nemec et al., 2017), which has drawbacks compared to regular, denaturing SDS-PAGE. On the other hand, GFP and FPs in general are known to be tightly folded and more resistant to denaturation than most proteins (Saeed and Ashraf, 2009; Ward, 2005). For example, a few studies exploited the ability of GFP to resist the adverse conditions of SDS-PAGE to run protein overexpression screens in *Escherichia coli* or other cell types (Aoki et al., 1996; Bird et al., 2015; Bomholt et al., 2013; Drew et al., 2006; Geertsma et al., 2008; Krasnoselska et al., 2021; Madani et al., 2021; Muller-Lucks et al., 2012; Newstead et al., 2007), or on recombinantly expressed or purified proteins *in vitro* (Aoki et al., 1996; Campbell et al., 2002; Donate-Macian et al., 2019; Koldenkova et al., 2015; Nakatani et al., 2019; Weinberger Ii and Lennon, 2021; Yanushevich et al., 2002).

The development and popularity of fluorescence imaging devices is likely to lead to a generalization of the use of FPs for direct visualization of in-gel fluorescence (IGF). Accordingly, a recent study used this method for easy detection of FP-tagged proteins in protein extracts of cell culture (Ruan et al., 2024).

In this study, we document that in-gel fluorescence (IGF) detection is a rapid and cost-effective alternative to immunoblotting to visualize FP-tagged proteins expressed at endogenous levels in cell extracts. IGF avoids the time-consuming process of transfer to a membrane, incubations with antibodies and washes, with substantial improvements of the cost of the experiment as this does not require primary nor secondary antibodies, membranes, or chemiluminescence substrates, and of the quality of the data (less background and higher dynamic range). We describe the conditions of use and limitations of various GFPs and RFPs for IGF, as well as sample preparation, imaging conditions, sensitivity and possible applications.

## Results

### The fluorescence of endogenously yeGFP-tagged proteins can be visualized from cell extracts in SDS-PAGE gel after migration

To know whether we can detect GFP fluorescence from endogenously expressed proteins in total protein extracts, we tagged two yeast genes (*BMH1*, encoding a 14-3-3 protein) or *HXK1*, encoding hexokinase 1) at their endogenous locus with yeGFP, a yeast codon-optimized GFP (Cormack et al., 1997) which also harbors two mutations (S65G, S72A - a.k.a. “GFPmut 3”, Cormack et al., 1996) selected to increase GFP fluorescence (ID on FPBase https://www.fpbase.org : A2OWC, Lambert, 2019). yeGFP is often used for fluorescent tagging as it is incorporated into a toolbox for PCR-based tagging of yeast genes at their endogenous genetic locus (Janke et al., 2004). Exponentially growing cells (5 OD equivalents) were lysed with glass beads at 4°C in 100 µL native lysis buffer (containing Tris-HCl, Triton X-100 and NaCl; see Material and Methods for more details). 4X Laemmli sample buffer (Bio-Rad, containing lithium dodecyl sulfate [LDS] and dithiothreitol [DTT]) was then added to the lysates (1X final), and samples were then incubated at 30°C or 95°C for 5 min, before loading on commercial precast TGX (Tris-Glycine eXtended) gels, and run in Tris-Glycine-SDS (TGS) buffer. After migration, the gel was imaged with gel-imaging devices whose excitation/emission wavelengths were compatible with the observation of green fluorescent proteins (ChemiDoc MP, Bio-Rad: 460-490/518-546 nm; and Typhoon, Cytiva: 488/505-545 nm) (**Figure S1**). The gels displayed a green fluorescent signal at the size of the tagged proteins, as revealed by immunoblotting with anti-GFP antibodies after transfer of the same gel to a membrane (**Figure 1A**). Thus, at least some GFP molecules remained fluorescent throughout sample preparation and gel migration process. Noteworthy, other lysis buffers were tested and also allowed the maintenance of GFP fluorescence (see Material and Methods for more details). On a side note, we observed that gel exposure to UV-light (such as during total protein labeling with a trihalo compound, eg. “Stain-Free” technology from Bio-Rad: 45 sec exposure at 300-400 nm) partially bleached the GFP fluorescence signal (**Figure S2**). Thus, this labeling should preferentially be performed after visualizing the GFP signal, especially if the signal is weak. Finally, as expected, GFP was no longer fluorescent when samples were denatured at 95°C instead of 30°C, or when extracts were prepared from cells precipitated with TCA in which proteins are fully denatured (**Figure 1A**).

**Figure 1.**
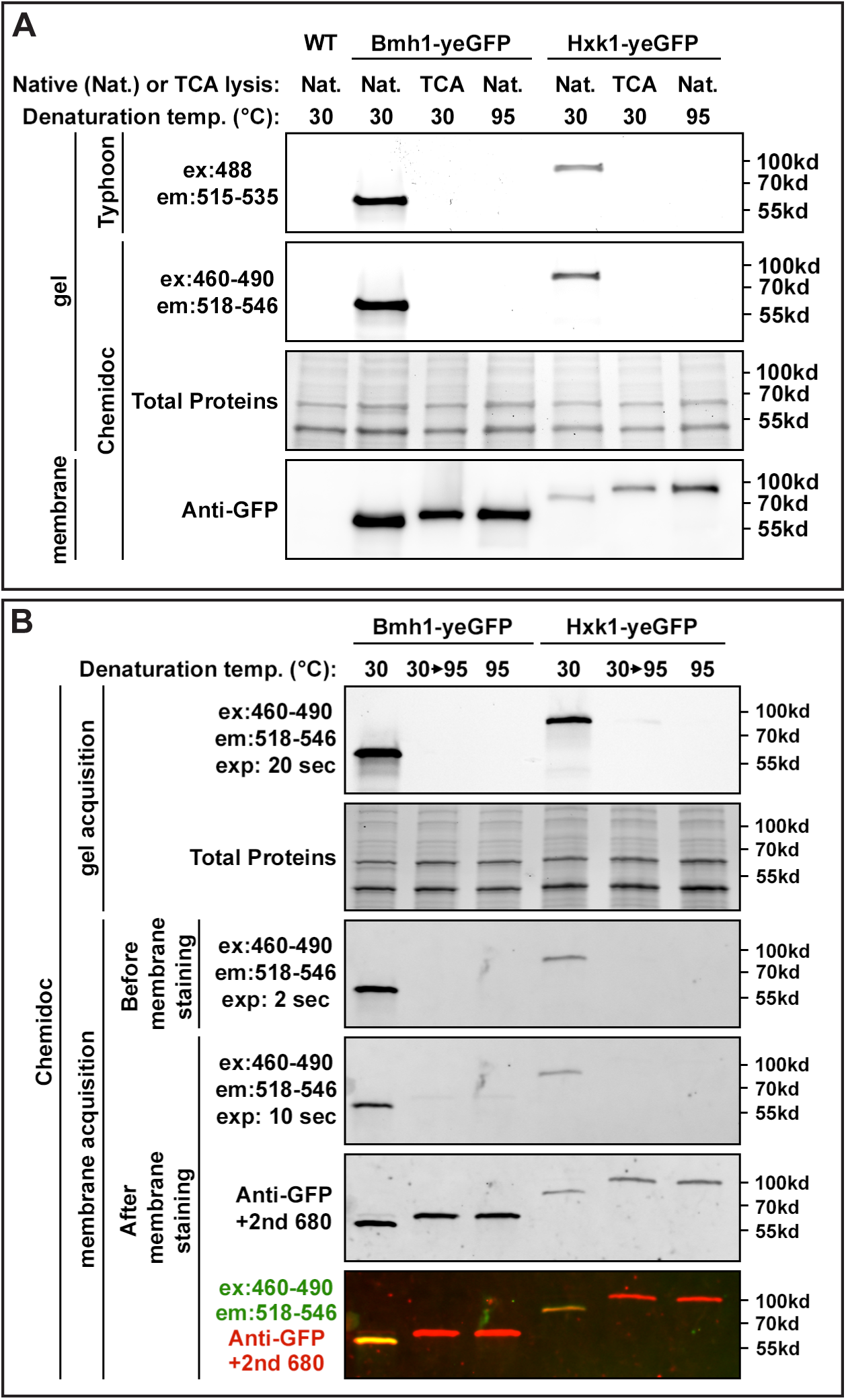
Detection of in-gel fluorescence from endogenously expressed proteins tagged with yeGFP. A. Control (WT) yeast or yeast expressing Bmh1-yeGFP, Hxk1-yeGFP were lysed in the indicated conditions (Nat.: native conditions; TCA: lysis in denaturing conditions with TCA), samples were resuspended in LDS sample buffer (1X final, Bio-Rad) and incubated for 5 min at the indicated temperatures. After migration on a commercial precast 4-20% TGX gels (Bio-Rad), gels were imaged for green fluorescence using a Typhoon and a Chemidoc MP, and total proteins were visualized by the stain-free technology on a Chemidoc MP. After transfer to a nitrocellulose membrane, proteins were immunoblotted with anti-GFP antibodies (Roche) and detected by chemiluminescence using anti-mouse antibodies coupled to HRP. B. Protein lysates were prepared as in A. Proteins were incubated at 30°C or 95°C for 5 min, with and without prior incubation at 30°C (5 min). After migration on a commercial precast 4-20%TGX gel (Bio-Rad), gels were imaged for green fluorescence using a Chemidoc MP, and total proteins were visualized by the stain-free technology on a Chemidoc MP. Proteins were then transferred to a nitrocellulose membrane and imaged again for yeGFP fluorescence, which was maintained during transfer. Proteins were then immunoblotted with anti-GFP antibodies (Roche) and anti-mouse antibodies coupled to Alexa Fluor 680. yeGFP fluorescence was then visualized again before detecting the anti-GFP antibodies by fluorescence on a Chemidoc MP.

Whereas these mild denaturing conditions allow the detection of in-gel fluorescence, two important points should be noted. First, the lower level of denaturation can impact on the migration of the GFP-tagged protein. At 30°C, Bmh1-yeGFP and Hxk1-yeGFP migrated faster than when samples were heated at 95°C or precipitated with TCA, as determined after transfer and immunoblotting with anti-GFP antibodies (**Figure 1A**). In-gel fluorescence detection systematically correlated with the presence of the fast-migrating band, suggesting it may correspond to a folded species of yeGFP, in agreement with previous observations (Aoki et al., 1996). To address this point, proteins were transferred onto a nitrocellulose membrane in classical conditions (liquid transfer), and immunoblotted with anti-GFP antibodies. Bmh1-yeGFP fluorescence signal could be detected on the nitrocellulose membrane even after transfer and incubation with antibodies, allowing the sequential observation of yeGFP fluorescence and antibody-based labeling during the same acquisition. This revealed that the yeGFP fluorescence signal precisely overlaid with the fast-migrating band detected with the anti-GFP antibody (**Figure 1B**). Thus, yeGFP remains mostly folded in these mild denaturation conditions allowing in-gel fluorescence observation, but can also cause a variation in the apparent molecular weight of yeGFP-tagged proteins as compared to denatured proteins.

A second aspect that should be considered when using low denaturing conditions is that various antibodies may recognize folded and denatured fluorescent proteins with different affinities. For instance, anti-GFP antibodies may display a greater affinity towards folded or unfolded GFP, leading to potential artifacts during quantification of the signal (**Figure S3A**). This could be circumvented by denaturing proteins directly on the membrane after transfer (**Figure S3B**, see Material and Methods).

### Comparison of GFP(S65T), yeGFP and EGFP for in-gel fluorescence visualization

Following the development of tools for tagging of yeast genes with various tags by homologous recombination (Longtine et al., 1998; Wach et al., 1997), Huh et al. generated a collection of yeast strains which is widely used by the community and in which most genes are individually tagged with GFP(S65T) (Huh et al., 2003) (FPBase ID: B6J33). To evaluate whether strains from this collection are suitable for in-gel fluorescence detection, we retrieved the Bmh1-GFP(S65T) strain and subjected it to the same analysis. A fluorescent signal was detected in the gel when samples were incubated at 30°C, but not 95°C (**Figure 2A**). When compared to the signal obtained when tagging the same protein with yeGFP, the fluorescent signal observed with GFP(S65T) was weaker, probably because GFP(S65T) was less robust and more prone to denaturation. Indeed, a large fraction of the Bmh1-GFP(S65T) protein pool was denatured even when incubated at 30°C prior to gel loading, as judged by the migration pattern revealed with anti-GFP antibodies and antibodies directed to the proteins of interest (**Figure 2A**). Thus, GFP(S65T) is subject to a partial denaturation even in conditions which otherwise preserve yeGFP fluorescence. This was confirmed when comparing the staining of Hxk1-GFP(S65T) with that of Hxk1-yeGFP (**Figure 2A**). In conclusion, fusions with GFP(S65T) (including those from the GFP collection, Huh et al., 2003) are not the best-suited to detect in-gel fluorescence in our conditions.

**Figure 2.**
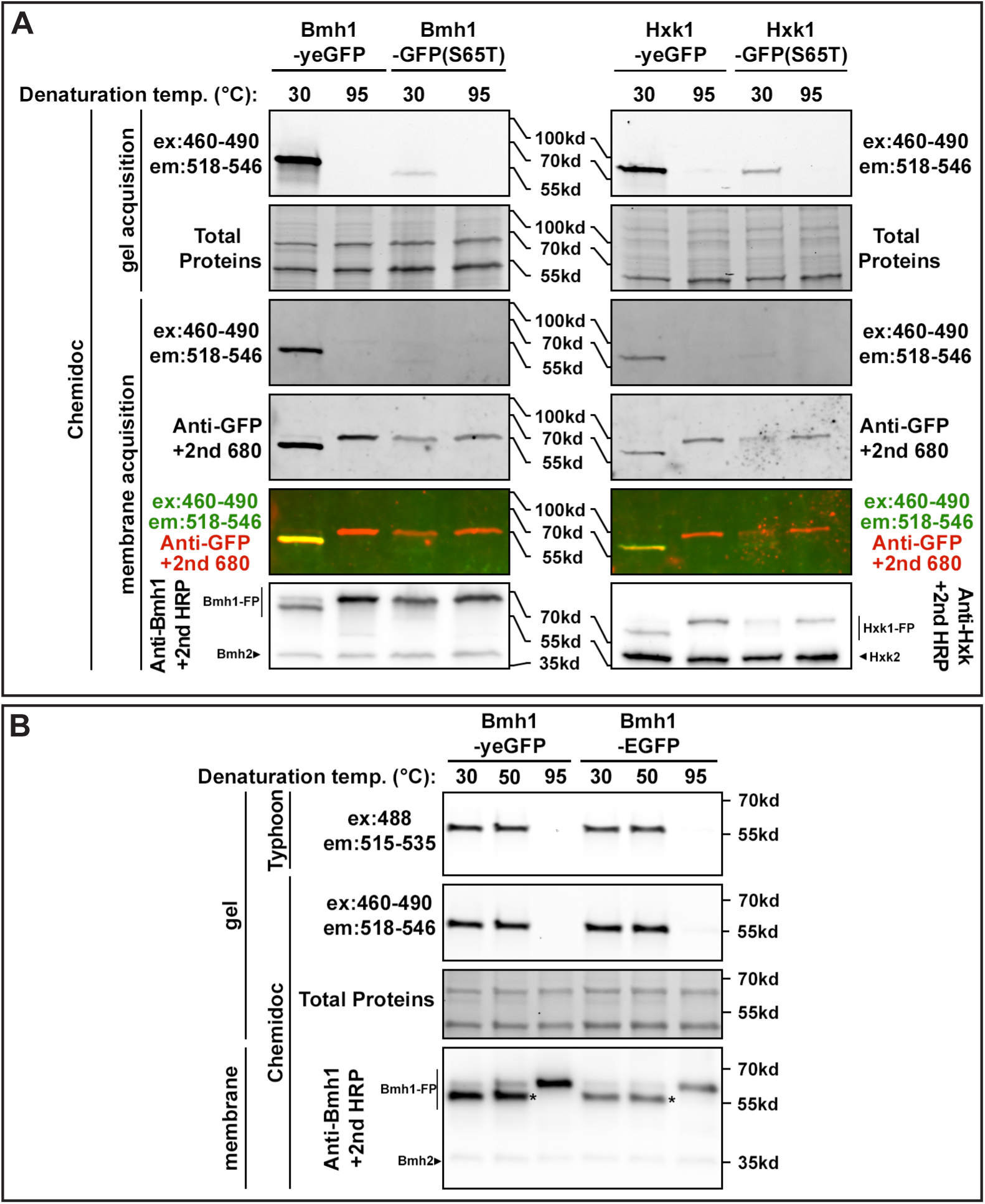
Comparison of in-gel fluorescence of yeGFP-, GFP(S65T)- and EGFP-tagged proteins. A. Yeast expressing Bmh1-yeGFP, Bmh1-GFP(S65T), Hxk1-yeGFP and Hxk1-GFP(S65T) were lysed in native conditions, samples were resuspended in LDS sample buffer and incubated for 5 min at the indicated temperatures. After migration on a commercial precast 4-20% TGX gels (Bio-Rad), gels were imaged for green fluorescence using a Chemidoc MP, and total proteins were visualized by the stain-free technology on a Chemidoc MP. Proteins were then transferred to a nitrocellulose membrane and imaged again for green fluorescence, which was maintained during transfer. Proteins were then immunoblotted with anti-GFP antibodies (Roche) and anti-mouse antibodies coupled to Alexa Fluor 680. Green fluorescence was then visualized again before detecting the anti-GFP antibodies by fluorescence. The membranes were then stripped and incubated with anti-Bmh1 or anti-Hxk1/2 antibodies and then with anti-rabbit antibodies coupled to HRP, and revealed by chemiluminescence on a Chemidoc MP. B. Yeast expressing Bmh1-yeGFP and Bmh1-EGFP were lysed in native conditions, samples were resuspended in LDS sample buffer and incubated for 5 min at the indicated temperatures. After migration on a commercial precast 4-20% TGX gel (Bio-Rad), gels were imaged for green fluorescence using a Typhoon or a Chemidoc MP, and total proteins were visualized by the stain-free technology on a Chemidoc MP. Proteins were then transferred to a nitrocellulose membrane and immunoblotted with anti-Bmh1 antibodies and then with anti-rabbit antibodies coupled to HRP, and revealed by chemiluminescence on a Chemidoc MP. * indicates the fluorescent species.

We also tested another commonly used GFP variant, EGFP, which derives from the original avGFP and carries mutations F64L and S65T (a.k.a. “GFPmut 1”, FPBase ID: R9NL8, Cormack et al., 1996). This variant is also commonly used for the tagging of yeast genes at their endogenous their availability in the pEGFP-N1/pEGFP-C1 plasmids for expression in mammalian cells. The signals obtained with EGFP were comparable to those obtained for yeGFP, making it appropriate for visualization of IGF (**Figure 2B**).

### In-gel fluorescence detection of proteins tagged with mNeonGreen and superfolder GFP

We next tested the suitability of more recently developed green-fluorescent proteins for IGF detection (**Figure 3**). For comparative purposes, the same protein that we used for the initial variants tested, Bmh1, was tagged with the brighter dLanYFP-derivative, mNeonGreen (FPBase ID: ZRKRV, Shaner et al., 2013) or with superfolder GFP (sfGFP), a tightly folded GFP which is even more resistant to denaturation than traditional GFP (FPBase ID: B4SOW, Pedelacq et al., 2006).

**Figure 3.**
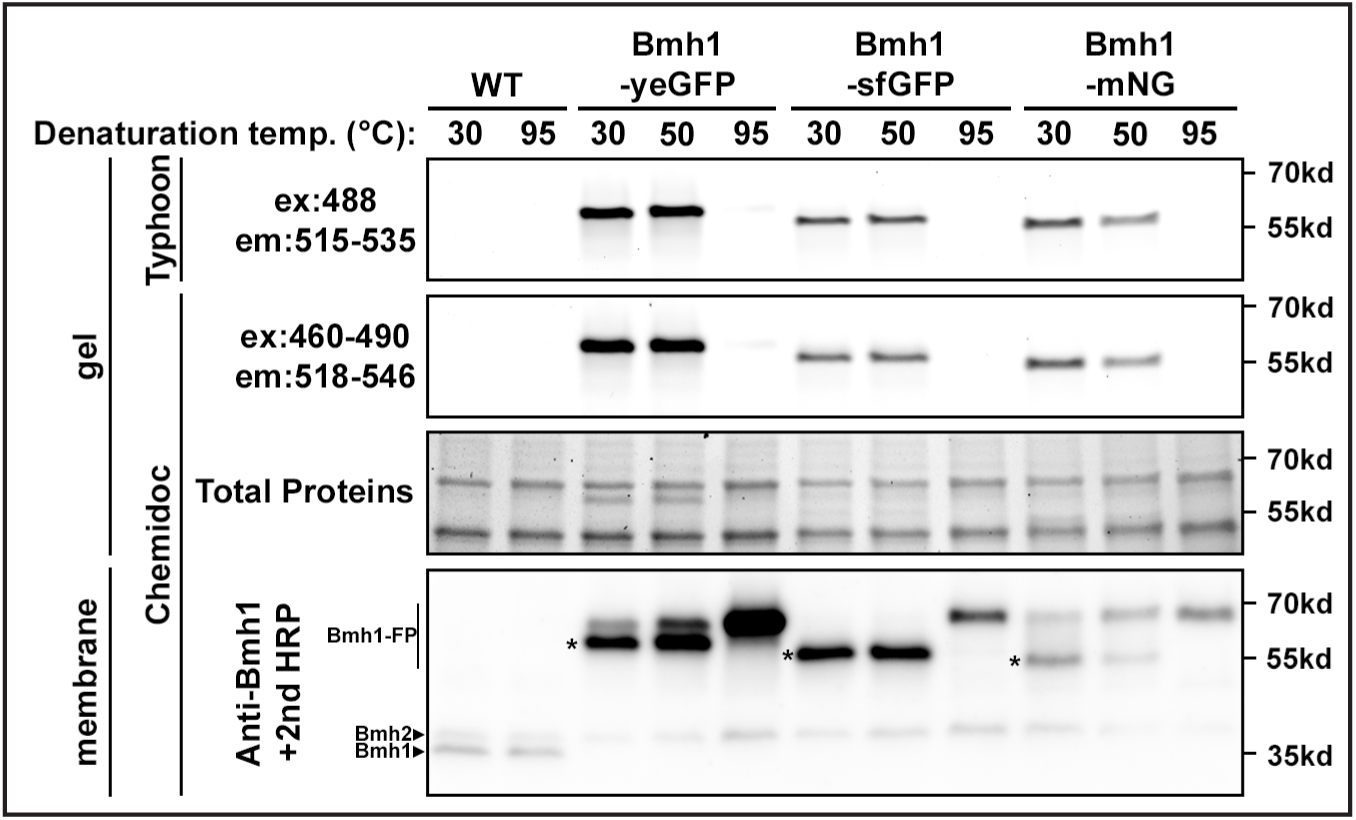
Comparison of in-gel fluorescence of yeGFP-, sfGFP- and mNeonGreen-tagged proteins. Yeast expressing Bmh1-yeGFP, Bmh1-sfGFP or Bmh1-mNeonGreen were lysed in native conditions, samples were resuspended in LDS sample buffer and incubated for 5 min at the indicated temperatures. After migration on a commercial precast 4-20%TGX gel (Bio-Rad), gels were imaged for green fluorescence using a Typhoon or a Chemidoc MP, and total proteins were visualized by the stain-free technology on a Chemidoc MP. Proteins were then transferred to a nitrocellulose membrane and immunoblotted with anti-Bmh1 antibodies and then with anti-rabbit antibodies coupled to HRP, and revealed by chemiluminescence on a Chemidoc MP. * indicates the fluorescent species.

In all cases, a fluorescent signal was detected in the gel when samples were incubated at 30°C prior to loading (**Figure 3**). Various fluorescence intensities were observed, with the strongest signal obtained for Bmh1-yeGFP. However, these variations could be due to (i) the spectral properties of these fluorescent proteins which may not appropriately match the imaging systems used for their detection, (ii) the ability of these fluorescent proteins to endure the sample preparation conditions and SDS-PAGE protocol, and (iii) an effect of the tag on the expression level of the tagged protein (**Figure 3**). Indeed, in our experiments, Bmh1-yeGFP was more highly expressed than Bmh1-mNeonGreen or Bmh1-sfGFP, as determined by immunoblotting these proteins with anti-Bmh1 antibodies. We hypothesized that this lower level of expression might be due to the fact that only Bmh1-yeGFP is codon-optimized for expression in yeast. However, optimizing codon usage for mNeonGreen expression in yeast (ymNeonGreen) neither led to an increase in protein expression nor in IGF signal (**Figure S4**). This hypothesis remains open for sfGFP, as we did not test optimization of codon usage for this FP.

The loss of IGF signal upon denaturation at 95°C was accompanied by a change in the migration pattern as revealed by western blotting using anti-Bmh1 antibodies, with only one band present at a higher apparent molecular weight, corresponding to the non-fluorescent species. An intermediate situation was observed upon denaturation at 50°C, with a decrease in Bmh1-mNeonGreen IGF, unlike what was observed for Bmh1-yeGFP or sfGFP (**Figure 3**). Accordingly, by immunoblotting, treatment of samples at 50°C led to an almost full conversion of Bmh1-mNeonGreen from the native to the denatured form, which was less pronounced for Bmh1-yeGFP. Thus, mNeonGreen is amenable to IGF detection but is less heat-stable than other green FPs. In contrast, Bmh1-sfGFP migration was unaffected at 50°C compared to 30°C, highlighting the stronger heat-stability of this fluorescent variant. Overall, we conclude that sfGFP and EGFP (see **Figure 2B**) are more resistant to heat-induced denaturation than yeGFP, GFP(S65T) and mNeonGreen.

These observations led us to quantify the heat-stability of fluorescence of the most robust green FPs, which could be helpful for applications in which samples must be incubated at temperatures (**Figure 4**). In-gel fluorescence of Bmh1-sfGFP was unaltered until reaching 59°C, whereas Bmh1-yeGFP fluorescence decreased starting at 54°C (**Figure 4A, C-D**). The temperature at which 50% of the signal is still present was 67°C for sfGFP and 59°C for yeGFP. Interestingly, EGFP behaved similarly to sfGFP (**Figure 4B, D**). The only additional gain of tagging with sfGFP over the widely used EGFP is that it migrates almost exclusively as a single band at low denaturing temperatures (30°C). Again, denaturation of all green FP-tagged Bmh1 was accompanied by a decrease in its mobility on gels, which could clearly be visualized using anti-Bmh1 antibodies (**Figure 4, A-C**).

**Figure 4.**
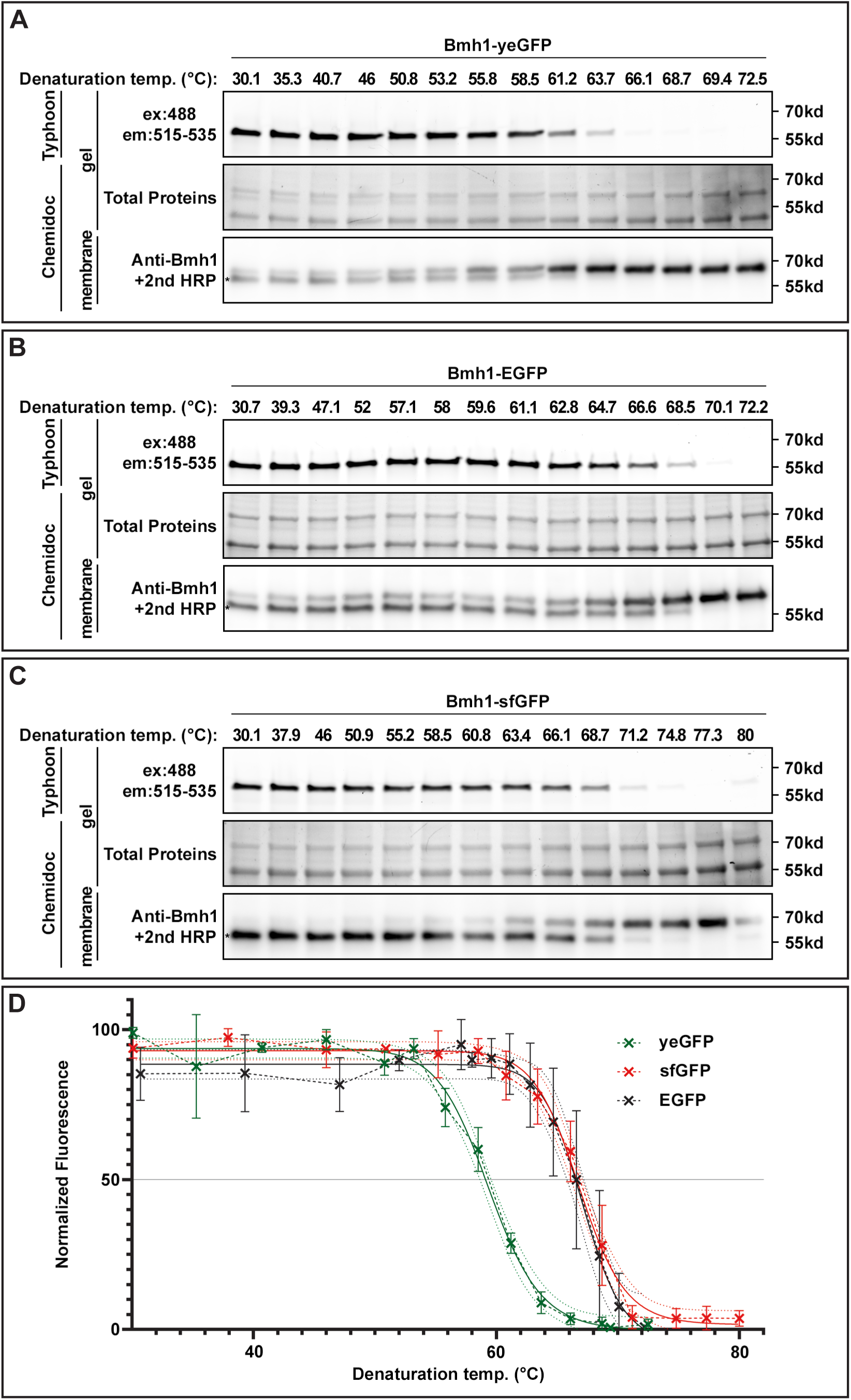
Temperature-sensitivity of in-gel fluorescence of yeGFP-, EGFP- and sfGFP-tagged proteins. A. Yeast expressing Bmh1-yeGFP were lysed in native conditions, samples were resuspended in LDS sample buffer and incubated for 5 min at the indicated temperatures in a gradient thermocycler (note that the temperature range is different for sfGFP). After migration on a commercial precast 4-20%TGX gel (Bio-Rad), gels were imaged for green fluorescence using a Typhoon, and total proteins were visualized by the stain-free technology on a Chemidoc MP. Proteins were then transferred to a nitrocellulose membrane and immunoblotted with anti-Bmh1 antibodies and then with anti-rabbit antibodies coupled to HRP, and revealed by chemiluminescence on a Chemidoc MP. B. Same as A. using yeast expressing Bmh1-EGFP. C. Same as A. using yeast expressing Bmh1-sfGFP. D. Quantification of the green fluorescence signal as a function of the denaturation temperature for various green FP-tagged Bmh1 (n=3; ± SD). Solid line: sigmoidal fit of the data (GraphPad Prism), dotted line: 95% confidence interval of the fit.

Altogether, these results allow us to reach the following conclusions. First, Bmh1-yeGFP and Bmh1-EGFP give a strong IGF signal that, in our experiments, could be attributed to a higher expression level coupled to a relatively high resistance to denaturation, especially in the case of EGFP. Indeed, EGFP was as resistant as sfGFP to heat-induced denaturation in our conditions. Second, despite being more sensitive to the conditions of extraction and imaging, mNeonGreen in-gel fluorescence can still be used at low denaturation temperatures, which can avoid the purchase of specific antibodies – indeed, mNeonGreen shares only 25% identity with the traditional avGFP and is not recognized by anti-GFP antibodies. Finally, care should be taken regarding the expression level of the tagged protein which, despite tagging at the endogenous genetic locus, can lead to strong variations in expression depending on which FP was used for tagging (see **Figure 3**).

### In-gel fluorescence of red and orange fluorescent proteins

We aimed at expanding our observations to other, non-green fluorescent proteins. We could not test blue-fluorescent proteins because none of the commonly-used imaging devices (Chemidoc MP and Typhoon) provide filters compatible with the excitation/emission spectra of blue-fluorescent proteins. However, these experiments were possible for red-fluorescent proteins. We tagged Bmh1 with the widely used fluorescent protein mCherry, a monomeric DsRed derivative with many advantages such as high photostability, faster maturation and high pH-stability (FPBase ID: ZERB6, Shaner et al., 2004). We also fused Bmh1 with mRuby2, a bright derivative of eqFP611 (FPBase ID: 8MJ78, Lam et al., 2012), and with TagRFP-T, which derives from eqFP578 (Merzlyak et al., 2007) with enhanced photostability (FPBase ID: LF3LJ, Shaner et al., 2008). Finally, we also tested fusion of Bmh1 with mKO-κ, a derivative of the monomeric orange fluorescent protein mKO (Karasawa et al., 2004) further mutagenized for faster maturation (FPBase ID: HMK8R, Tsutsui et al., 2008) because of its spectral properties that make it more appropriate for detection by the Chemidoc MP (**Figure S5**). Protein extracts were run on SDS-PAGE and in-gel fluorescence was observed at excitation/emission wavelengths of 532/550-800nm (Typhoon) or 520-545/577-613nm (ChemiDoc MP). All samples gave a fluorescent signal in gel (**Figure 5A**). Fluorescence signals were globally stronger when using a long-pass filter on the Typhoon (550-800nm), which collects more fluorescence signals than the narrower filter present on the Chemidoc MP (577-613nm) (**Figure S5**). The strongest fluorescence signals were obtained for Bmh1 mKO-κ and Bmh1-mCherry, whereas migration patterns comparable to those obtained for green fluorescent proteins: a faster-migrating band corresponded to the fluorescent species, whereas the slower migrating form corresponded to the denatured pool. The ratio of fluorescent over denatured proteins was higher for mCherry, although a minor denatured pool was already present in mild denaturing conditions. Notably, the fluorescent pool of Bmh1-mRuby2 and Bmh1 mKO-κ migrated much faster than any other FP-tagged Bmh1 protein, with a decrease of about 15 kDa in apparent MW compared to the denatured version. Bmh1-mKO-κ was the best detected red fluorescent protein on the Chemidoc MP (**Figure 5A**), likely because of its spectral properties (**Figure S5**) which were more appropriate for detection on this device; however, a strong signal was also observed for Bmh1-mCh. Altogether, our conclusions are that (i) mild denaturation conditions are compatible with IGF of orange and red FPs, with mCherry and mKO-κ being the best tags in our hands, with a stronger signal and resistance to heat-induced denaturation; (ii) IGF allows the visualization of proteins tagged with orange and red FPs originating from various organisms without the need to purchase specific antibodies; (iii) tagging of proteins with orange and red FPs can lead to significant changes in the apparent molecular weight of the fusion proteins in mild denaturing conditions.

**Figure 5.**
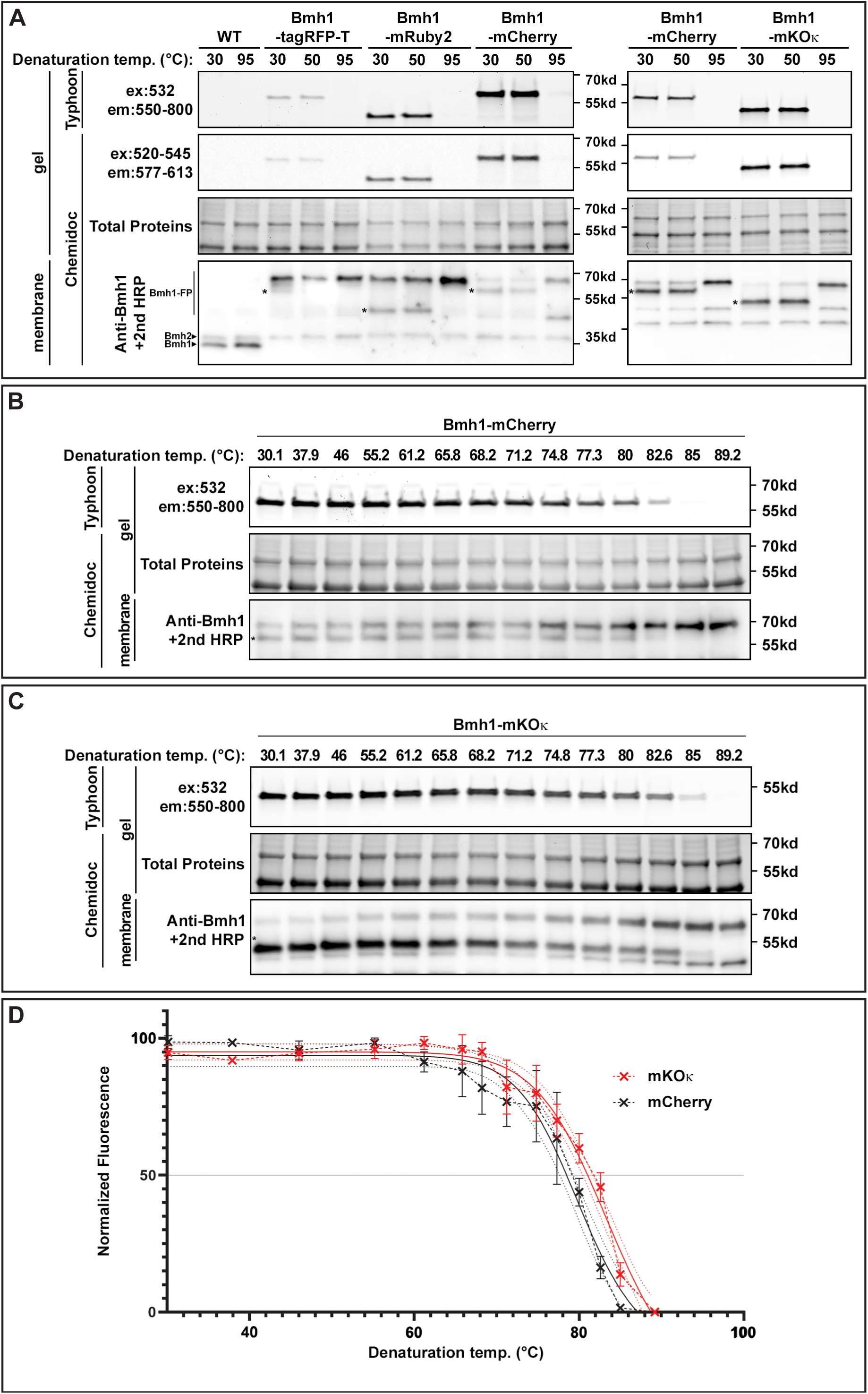
Comparison of in-gel fluorescence of tagRFP-T-, mRuby2-, mCherry- and mKO-κ-tagged proteins. A. Yeast expressing Bmh1-tagRFP-T, Bmh1-mRuby2, Bmh1-mCherry or Bmh1-mKO-κ were lysed in native conditions, samples were resuspended in LDS sample buffer and incubated for 5 min at the indicated temperatures. After migration on a commercial precast 4-20%TGX gel (Bio-Rad), gels were imaged for red fluorescence using a Typhoon or a Chemidoc MP, and total proteins were visualized by the stain-free technology on a Chemidoc MP. Proteins were then transferred to a nitrocellulose membrane and immunoblotted with anti-Bmh1 antibodies and then with anti-rabbit antibodies coupled to HRP, and revealed by chemiluminescence on a Chemidoc MP. * indicates the fluorescent species. B. Yeast expressing Bmh1-mCherry were lysed in native conditions, samples were resuspended in LDS sample buffer and incubated for 5 min at the indicated temperatures in a gradient thermocycler. After migration on a commercial precast 4-20%TGX gel (Bio-Rad), gels were imaged for red fluorescence using a Typhoon, and total proteins were visualized by the stain-free technology on a Chemidoc. Proteins were then transferred to a nitrocellulose membrane and immunoblotted with anti-Bmh1 antibodies and then with anti-rabbit antibodies coupled to HRP, and revealed by chemiluminescence on a Chemidoc MP. C. Same as (B) on lysates of yeast expressing Bmh1-mKO-κ. D. Quantification of the fluorescence signal as a function of the denaturation temperature for mCherry-tagged and mKO-κ-tagged Bmh1 (n=3; ±SD). Solid line: sigmoidal fit of the data (GraphPad Prism), dotted line: 95% confidence interval of the fit.

Because of the higher signal intensity provided by mKO-κ and mCherry, and the fact that the latter is a widely used FP, we then evaluated the resistance of Bmh1-mCh and Bmh1-mKO-κ to temperature-induced denaturation (**Figure 5, B-C**). Both mKO-κ and mCherry fluorescence was maintained at up to 60°C, with 50% of the signal still being present at 79°C (for mCherry) or 82°C (for mKO-κ). Thus, of all FPs tested (including green FPs), mCherry and mKO-κ are the most resistant to denaturation and are therefore perfectly appropriate for IGF signal detection.

### Linearity of the signal and sensitivity of IGF compared to antibody-based detection

Having established the capacity to detect fluorescence of numerous FPs in gels, we then turned to studying the linearity of the signal and the sensitivity of this approach compared to the traditional immunoblotting approach. When serial dilutions of Bmh1-yeGFP samples were initially assayed, we discovered that the simple dilution of the protein lysate in Laemmli buffer (containing SDS and DTT) was sufficient to promote FP denaturation, as determined by examining the migration profile using anti–Bmh1 antibodies. This was likely a consequence of an increase in the detergent/protein ratio, as previously observed (Xu and Keiderling, 2004) (**Figure S6A**). To circumvent this problem, we repeated the experiment using a non-denaturing sample buffer for dilutions of the sample (Laemmli buffer without LDS). Under these conditions, protein fluorescence was maintained in spite of the dilution (**Figure 6**).

**Figure 6.**
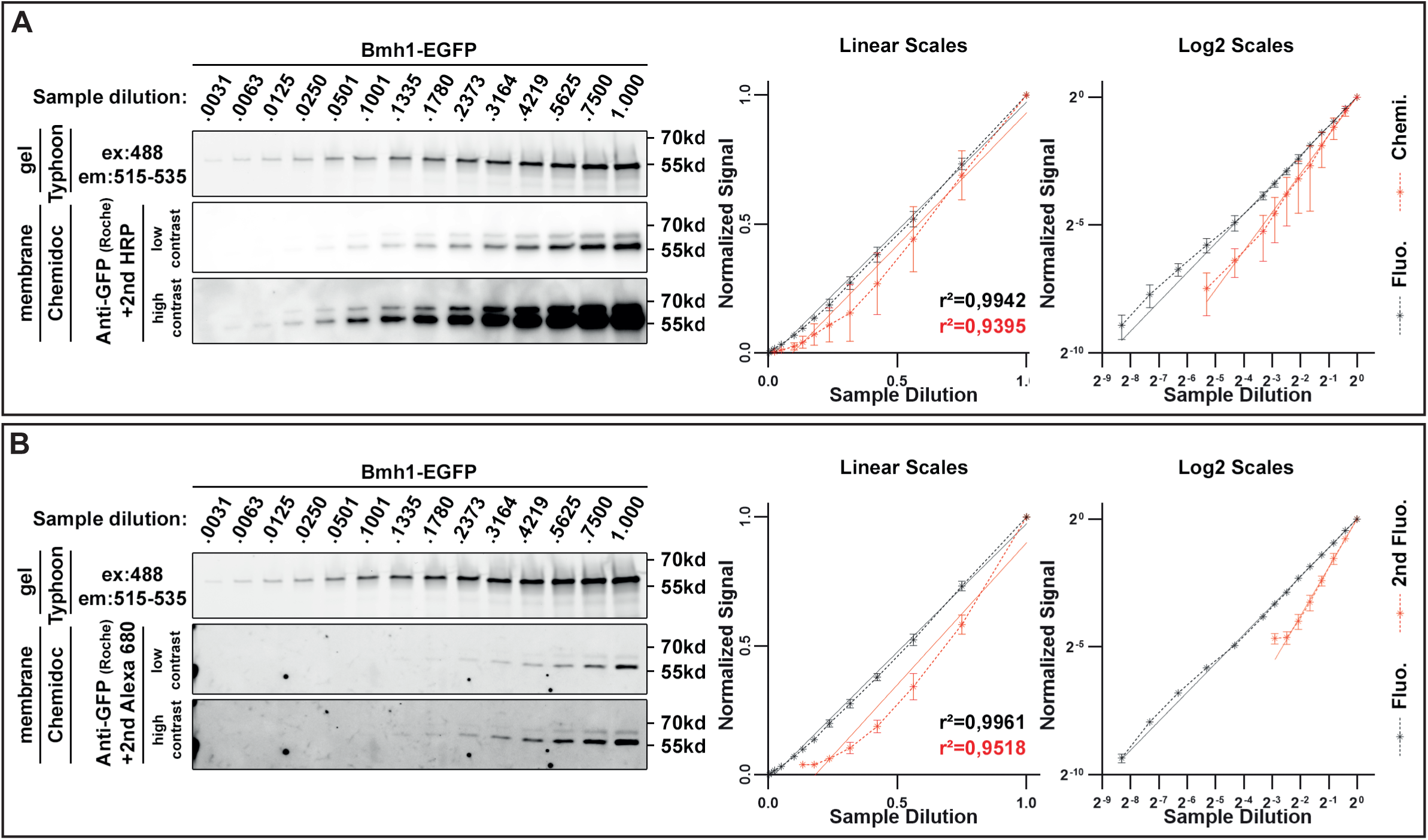
Sensitivity and linearity of in-gel fluorescence detection of EGFP. A. *Left.* Yeast expressing Bmh1-EGFP were lysed in native conditions, samples were resuspended in LDS sample buffer and incubated for 5 min at 30°C. Samples were serially diluted (right to left) into sample buffer without LDS (see Material and Methods), and loaded onto on a commercial precast 4-20%TGX gel (Bio-Rad). After migration, gels were imaged for green fluorescence using a Typhoon, and proteins were then transferred to a nitrocellulose membrane and immunoblotted with anti-Bmh1 antibodies and then with anti-rabbit antibodies coupled to HRP, and revealed by chemiluminescence on a Chemidoc MP. *Right.* Quantification of the signals obtained for green fluorescence (black) and chemiluminescence (red) as a function of sample dilution. Linear-scaled and log-plot scaled graphs are shown. The regression coefficient of a linear fitting is indicated. B. *Left.* Same experiment as in A, but using anti-GFP antibodies and anti-mouse antibodies coupled to HRP to reveal proteins by chemiluminescence. *Right.* Quantification of the signals obtained for fluorescence (black) and chemiluminescence (red) as a function of sample dilution. Linear-scaled and log-plot scaled graphs are shown. The regression coefficient of a linear fitting is indicated.

In-gel fluorescence was compared to the signal obtained using primary anti-GFP antibodies followed by secondary antibodies coupled to either horseradish peroxidase (HRP) for chemiluminescent detection (**Figure 6A**) or to a fluorescent dye (**Figure 6B**). Overall, IGF detection conditions. Overall, we found the IGF signal to more closely follow a linear plot than antibody-based signals, over a range of 320-fold (**Figure 6B**). The linearity of signal intensity as a function of protein dilution was confirmed for all FPs we tested, including mCherry and mKO-κ (**Figure S7**). Thus, we conclude that IGF detection is more reliable and sensitive than antibody-based detection.

Our observations above (**Figure S6A**) suggested that sample dilution in Laemmli buffer compromises the integrity of GFP. Although we observed that this could be circumvented by the use of detergent-free Laemmli buffer (**Figure 6**), we sought to find conditions in which sample dilution would maintain FP integrity despite the presence of detergent (**Figure S6B**). First, we observed that dilution of the protein lysate in native lysis buffer (Tris-HCl 100 mM pH 8.0, NaCl 0.15 M, Glycerol 5%, Triton X-100 0.5 %) instead of water (**Figure S6B**) led to a protection of fluorescence, as determined both by IGF detection and the observation of native/denatured species by immunoblotting (**Figure S6C**). This result could partially be attributed to the presence of NaCl in the native lysis buffer, which preserved fluorescence in itself (**Figure S6C**). Moreover, increasing the pH to 8.0 or above was sufficient to protect fluorescence at low protein concentrations (**Figure S6D**), allowing IGF detection even for diluted protein samples. We conclude that increasing the sample pH to pH 8.0 protects GFP denaturation in diluted samples.

### IGF detection on various endogenously-tagged proteins from yeast

We then aimed at detecting various proteins expressed at their endogenous level to address several questions. For these experiments, proteins were tagged with EGFP given its superior performances (see **Figure 2B**).

First, because Bmh1 and Hxk1 are rather abundant proteins (65,000 and 41,000 copies/cell, respectively; Ho et al., 2018), we determined IGF signals for proteins of lower abundance. IGF signal were obtained from proteins of abundance ranging from 736 to 65471 copies/cell (Ho et al., 2018) and originating from various subcellular locations (cytosol, nucleus, mitochondria, microtubules, plasma membrane, ER membrane) (**Figure 7A**). All proteins gave a signal at the expected size (**Figure 7B**), suggesting that IGF is applicable to proteins expressed at a low level. IGF signal intensity was then correlated with the published abundance originating from multiple studies (Ho et al., 2018) (**Figure 7C**). This revealed a good correlation between our IGF quantitation and these data, regardless of the type of experiments used for quantitation (quantitative proteomics, western blotting or confocal microscopy) (**Figure S8A**). It was also clear that in our conditions, IGF led to an overall better quality of detection than antibody-based chemiluminescence (**Figure S8B**).

**Figure 7.**
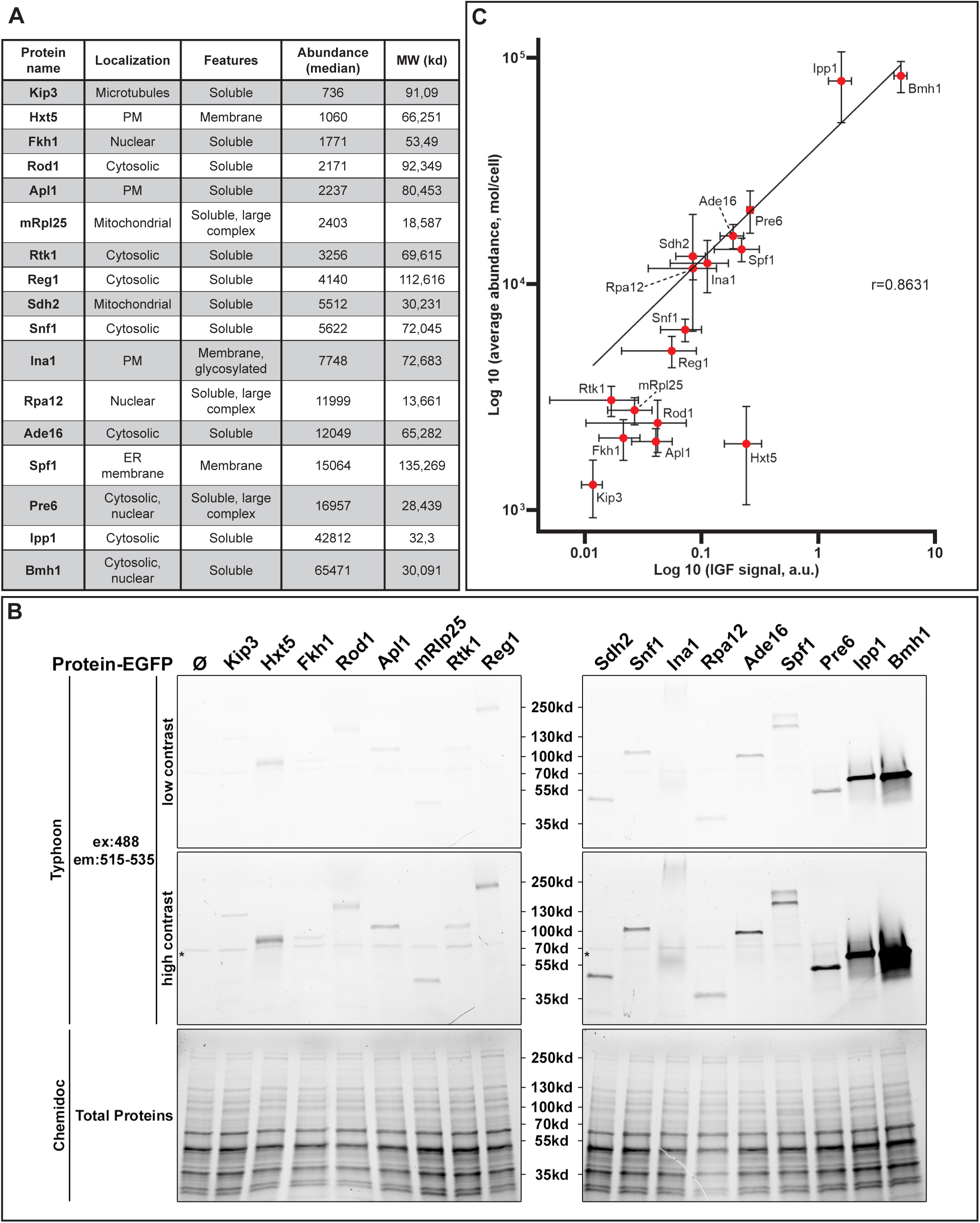
Detection of endogenous yeast EGFP-tagged proteins with various expression levels and correlation with published abundances. A. Proteins tagged with EGFP for IGF detection. Abundance (median) is according to Ho et al., 2018 (Ho et al., 2018) and available at www.yeastgenome.org. PM: plasma membrane, ER: endoplasmic reticulum. B. Protein lysates prepared in native conditions from yeast expressing the indicated proteins fused to EGFP. Samples were resuspended in LDS sample buffer and incubated for 5 min at the 30°C and loaded onto on a commercial precast 4-20%TGX gel (Bio-Rad). After migration, gels were imaged for green fluorescence using a Typhoon, and total proteins were visualized by the stain-free technology on a Chemidoc MP. A representative gel is shown.

Second, to check whether our conditions were sufficient to disrupt protein-protein interactions at the low denaturation temperatures we use, we tested proteins engaged in large protein complexes. Among the proteins we tested, several belonged to large complexes, such as RNA polymerase I (Rpa12), proteasome (Pre6), or mitochondrial ribosome (mRpl25). All of these migrated at the expected size (**Figure 7B**), with only one band being detected, suggesting disruption

Third, we also looked at proteins from various subcellular compartments as well as membrane proteins. The plasma membrane glucose transporter Hxt5 and the ER membrane-localized P-type ATPase Spf1 migrated at the appropriate size, although a doublet was observed for Spf1 as previously reported (Corradi et al., 2012; Hovsepian et al., 2017). The plasma membrane protein Ina1 also migrated as a diffuse band at the expected size given its high glycosylation, as previously reported (Laussel et al., 2022).

Altogether, our results show that IGF allows to visualize soluble and membrane proteins from various compartments, even when expressed at a modest level, and that the mild denaturation conditions are sufficient to denature high-order protein complexes.

### Use of IGF for the detection of FP-tagged proteins interaction by co-immunoprecipitation and compatibility with fluorescent SNAP-tagging

We sought to exploit the power of IGF detection in a situation in which two fluorescent proteins must be detected. Protein-protein interactions can be studied by the co-expression of tagged proteins and immunoprecipitation of one partner to reveal the presence of the potential partner in the immunoprecipitate. As a case study, we studied the interaction between the yeast 14-3-3 proteins Bmh1 and Bmh2, which are known to heterodimerize *in vivo* (Chaudhri et al., 2003). Lysates of yeast expressing Bmh1-yeGFP and Bmh2-ymCh (a yeast codon–optimized version of mCherry, see Material and Methods) were subjected to immunoprecipitation using nanobodies-based traps. Similar to our observations on diluted samples (section above), we realized that immunoprecipitates of FP proteins were denatured in the presence of Laemmli buffer, likely because of the low protein concentration in these samples. Based on our findings above (**Figure S6**), immunoprecipitates were resuspended in Laemmli buffer at pH 8.0 before loading, and samples were migrated on SDS-PAGE and imaged for IGF. In both cases (GFP-trap: **Figure 8A**; RFP-trap: **Figure 8B**), IGF detection allowed visualization of both the bait and the prey on the same gel after exposure to the appropriate wavelengths, within minutes after uncasting the gel. One limitation was a bleed-through of intense green fluorescence signals in the red channel, which was notably observed using the Chemidoc (**Figure S9**) but also using the Typhoon to a limited extent (**Figure 8A**). Thus, when looking at proteins tagged in both channels, the size of the green fluorescence-tagged protein should be different from that of the mCherry-tagged protein, and appropriate negative controls (samples without the GFP-tagged protein) should be included to determine the specificity of the signal.

**Figure 8.**
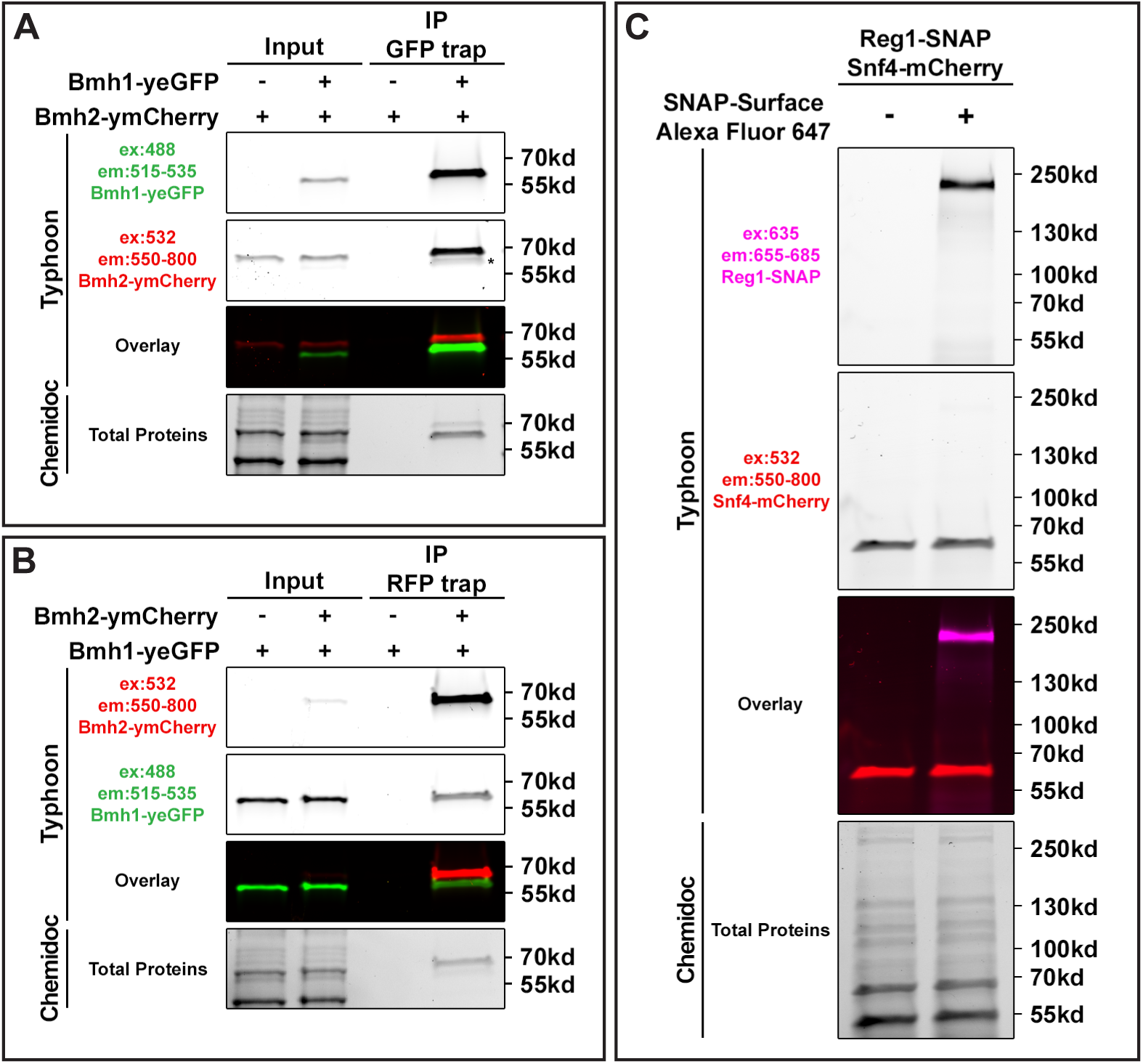
IGF detection in the context of co-immunoprecipitation or SNAP-tagging. A. Protein lysates prepared in native conditions with native IP lysis buffer from yeast expressing Bmh2-ymCherry (a yeast codon-optimized mCherry) with or without the co-expression of Bmh1-yeGFP were subjected to co-immunoprecipitation using GFP-trap beads. Immunoprecipitates were incubated at 50°C for 20 min in 1X Laemmli sample buffer at pH 8.0 which allows to maintain protein fluorescence at low protein concentrations. Input samples and immunoprecipitates (IP) are shown. After migration on a commercial precast 4-20%TGX gel (Bio-Rad), gels were imaged for green and red fluorescence using a Typhoon, and total proteins were visualized by the stain-free technology on a Chemidoc MP. * indicates bleed-through of the green fluorescence into the red channel. B. Protein lysates prepared in native conditions from yeast expressing Bmh1-yeGFP with or without the co-expression of Bmh2-ymCh were subjected to co-immunoprecipitation using RFP-trap beads and treated as in A. Input samples and immunoprecipitates (IP) are shown. After migration on a commercial precast 4-20%TGX gel (Bio-Rad), gels were imaged for green and red fluorescence using a Typhoon, and total proteins were visualized by the stain-free technology on a Chemidoc MP. C. Protein lysates prepared in native conditions with native IP lysis buffer from yeast expressing Reg1-SNAP and Snf4-mCherry, and were treated or not with SNAP-surface Alexa Fluor 647. After migration on a commercial precast 4-20%TGX gel (Bio-Rad), gels were imaged for red and far-red fluorescence using a Typhoon, and total proteins were visualized by the stain-free technology on a Chemidoc MP.

Because IGF is a fluorescence-based method, we also examined its compatibility with other approaches such as SNAP-tagging which allows a direct fluorescence-based detection and quantification of tagged protein in gel after electrophoresis (Tirat et al., 2006). A strain expressing Reg1-SNAP, tagged at its endogenous locus, was transformed with a plasmid expressing an mCherry-tagged protein (Snf4) under the control of its endogenous promoter. Total extracts were made in native conditions and SNAP fluorescence labeling was made after lysis using the SNAP-surface Alexa Fluor 647 dye which fluoresces in the far-red channel. This allowed detection of both mCh-tagged proteins and SNAP-tagged Reg1 in the gel after migration (**Figure 8C**). Therefore, IGF is compatible with SNAP-tagging protocols, further extending its abilities for in-gel quantitation of expression of multiple proteins.

### Use of IGF in cell extracts from other organisms

Although IGF detection is based on endogenous FP fluorescence and thus should not be affected by the cellular context in which these proteins are expressed, we tested the use of this technique in other eukaryotic systems, including Drosophila cells or tissues (**Figure 9, A-B**), MDCK cells (**Figure 9C**) and *Caenorhabditis elegans* whole organisms (**Figure 9D**). Protein extracts were prepared from FP-expressing cells in native conditions, resuspended in Laemmli buffer and samples were loaded on SDS-PAGE gels (see Material and Methods). Overall, only minor adaptations to current protein extraction protocols were made, and in all cases, IGF was observed, indicating the applicability of IGF detection to the analysis of protein lysates from other organisms. In particular, detection of the EGFP-tagged plasma membrane transducer Smoothened (SMO), which is phosphorylated in response to Hedgehog (Jia et al., 2004) shows that IGF detection can be used to study membrane proteins and post-translational modifications. In the case of total extracts from *C. elegans* worms, we could detect GFP and mCherry fluorescence signals, however, we observed the presence of non-specific fluorescent signals in the green channel that may complicate the readout of IGF. First, autofluorescence of *C. elegans* animals was previously reported (Pincus et al., 2016), and using red fluorescent proteins would avoid this problem. Second, we propose additional ways to circumvent this problem. A band migrating just below the 70-kDa marker appeared in lysates from WT animals which did not express any FP construct (**Figure 9D**), but this band disappeared upon denaturation at 50°C. A fluorescent smear of unknown origin was also observed at around 250 kDa (**Figure S10A**), which resisted the denaturation at 50°C. However, this smear was not visible when running the samples on homemade gels compared to commercial, pre-cast gels (**Figure S10B**), probably because this fluorescent material did not migrate into the resolving gel. Whereas these observations provide ways to circumvent the non-specific green fluorescence observed in *C. elegans*, it remains that care should be taken when setting up IGF detection in other material, including the use of negative, non-FP-tagged extracts to ensure the specificity of the signal.

**Figure 9.**
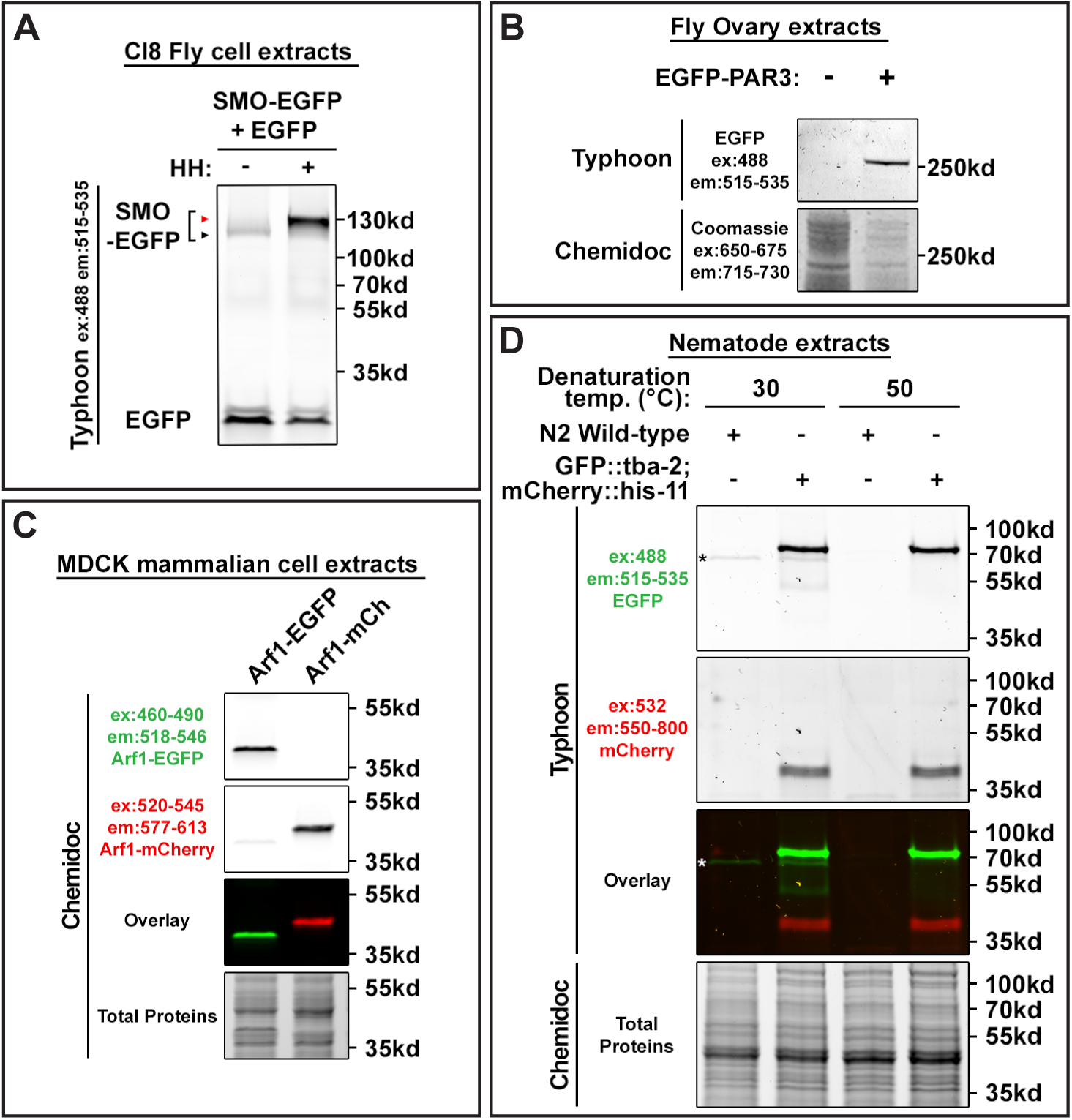
IGF detection in cells from various organisms. A. Drosophila wing-imaginal-disc cultured cells [Clone 8 (Cl-8) cells] were transfected with plasmids encoding the Hedgehog (HH) transducer Smoothened (SMO) fused to EGFP and EGFP alone. They were also transfected (+) or not (-) with a construct allowing the expression of HH. Cells were lysed in native conditions (see Material and Methods), lysates were mixed with 4X Laemmli sample buffer and incubated for 5 minutes at 25°C before loading on a homemade PAGE gel. IGF was detected on a Typhoon. SMO-EGFP is phosphorylated in the presence of HH, causing a slower migration (red arrowhead). B. Transgenic *Drosophila* flies expressing or not an EGFP-tagged version of the polarity protein PAR-3 were dissected and ovaries were lysed in native conditions (see Material and Methods). Samples were incubated for 5 min at 72°C in LDS sample buffer, loaded on a NuPage Bis-Tris gel (ThermoFisher) and IGF was detected on a Typhoon after migration. After fluorescence imaging, total proteins were stained with Coomassie stain and imaged on the infrared channel on a Chemidoc MP. C. MDCK cells expressing Arf1 fused to EGFP or mCherry were lysed in native conditions (see Material and Methods). Samples were incubated for 10 min at 50°C in LDS sample buffer and loaded on a gel. After migration on a commercial precast 4-20%TGX gel (Bio-Rad), gels were imaged for green and red fluorescence using a Chemidoc MP, and total proteins were visualized by the stain-free technology on a Chemidoc MP. Note that some bleed-through of green fluorescence is observed in the red channel (see also Figure 8A and **Figure S9**). D. Transgenic *C. elegans* expressing or not GFP-tagged alpha-tubulin and mCherry-tagged histone H2B were lysed in native conditions (see Material and Methods) by sonication. Samples were incubated for 5 min at 30°C or 50°C in Laemmli sample buffer (pH 8.0). After migration on a commercial precast 4-20%TGX gel (Bio-Rad), gels were imaged for green and red fluorescence using a Typhoon, and total proteins were visualized by the stain-free technology on a Chemidoc MP. * denotes a non-specific fluorescence protein present in extracts of WT *C. elegans* whose fluorescence disappears when heating the sample at 50°C.

## Discussion

Our study introduces in-gel fluorescence (IGF) detection as a rapid and cost-effective alternative to traditional immunoblotting for visualizing fluorescent protein (FPs) signals in cell extracts from yeast and other organisms, with minor adaptations of current protocols. The key points resulting from our observations and optimizations are summarized in **Figure 10**.

**Figure 10.**
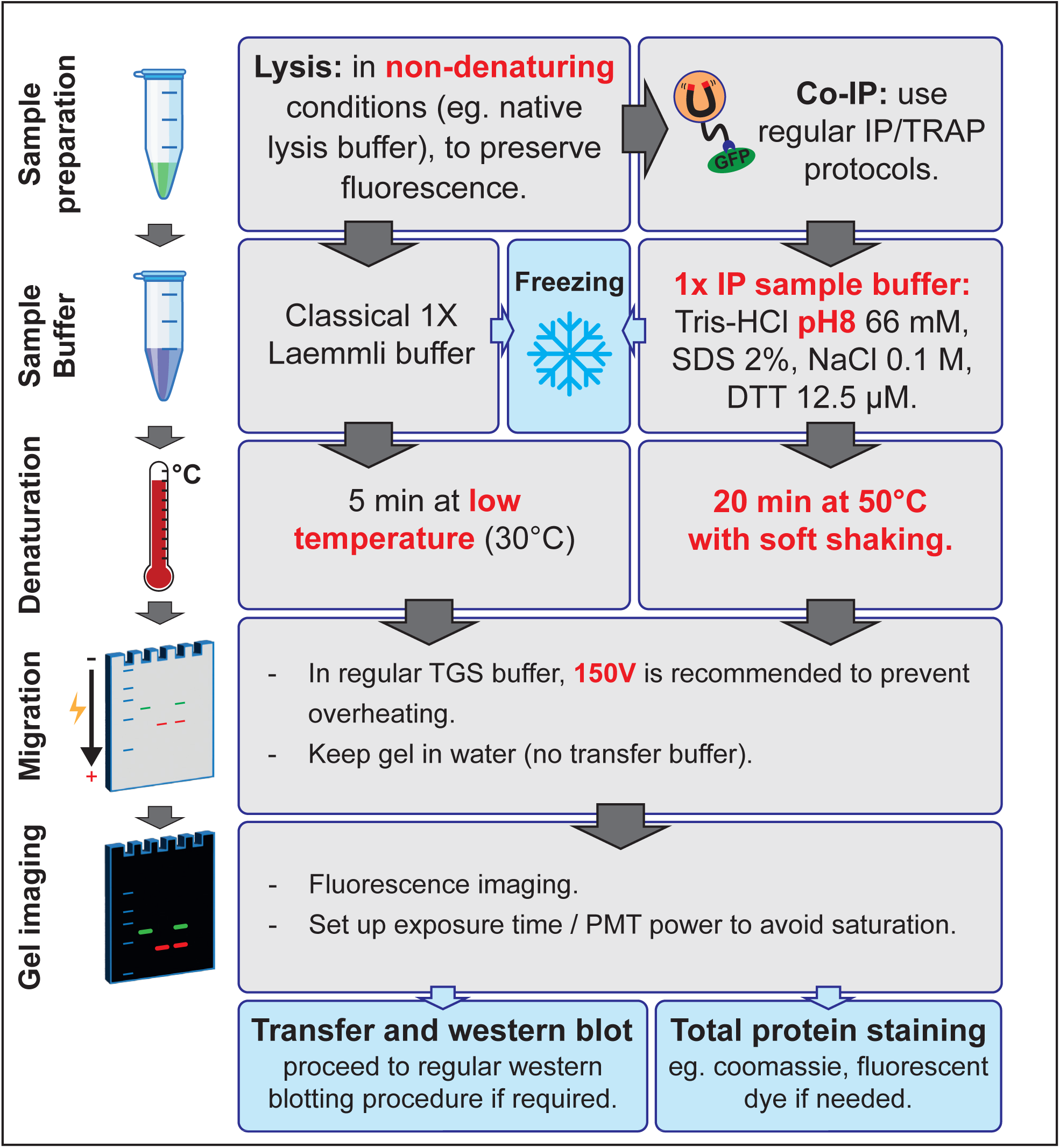
Overview of the workflow and technical considerations when using IGF detection to visualize proteins in gel. Overview of the IGF protocol. Lysis should be made in non-denaturing conditions to preserve endogenous fluorescence. Reagents tested include TX-100 (1%, Figure 9A), EDTA (10 mM, Figure 9C), DTT (2 mM, Figure 9C), antiprotease and antiphosphatase cocktails (Figure 9C), PMSF (0.5 mM), NaF (10 mM, Figure 8). Samples can be resuspended in classical Laemmli buffer containing SDS or LDS (12.5 µM DTT). Denaturation can be done preferentially for 5 min at 30°C, although longer times and higher temperatures were also tested throughout the study (**Figures 3, 4, 5**). For immunoprecipitates (Tris-HCl 66 mM pH 8.0, SDS 2%, NaCl 0.1 M, DTT 12.5 µM) to avoid denaturation occurring in diluted samples (**Figure S6B**). Samples can be frozen if needed (-20°C, - 80°C, liquid nitrogen, **Figure S2A**). Migration was generally done in TGX gels in regular TGS buffer, although NuPAGE Bis-Tris Gels were also used in MOPS buffer (Figure 9B). Migration at lower voltage than usual prevents overheating and FP denaturation. For faster migrations, the electrophoresis tank can be placed in ice-cold water, or pre-chilled buffer can be used. Note that Prestained protein ladders are fluorescent (**Figure S10**). Use unstained ladder or load a few microliters of prestained ladder. After uncasting, cut blue migration front to avoid fluorescence signal. Keep gel in water (no transfer buffer) and handle with clean gloves to avoid stains. Fluorescence imaging should be preferentially done before visualizing total proteins with UVs (“Stain-free”) (**Figure S2B**). Note that green fluorescence may bleed through in the red channel (Figures 8A**, S9**). Total protein staining can also be performed by Coomassie staining, which can be quantified in the near-infrared (**Figure S9B**) or with silver nitrate. Transfer can also be performed after fluorescence imaging, with maintenance of the FP fluorescence on the membrane (Figure 1) if no Ponceau staining was applied, and before addition of the chemiluminescent reagent as fluorescence is not maintained in hydrogen peroxide solution. Protein denaturation on membrane can increase antibody detection of low temperature-treated samples (**Figure S3B**).

Our work builds on the widespread use of FPs, particularly green and red fluorescent proteins and their derivatives, as powerful tools in cell biology. GFP, originally developed as a live-cell imaging marker, has since become key for various applications, including the monitoring of protein expression and as a proteomic tool. Immunoblotting has traditionally been the method of choice to visualize and quantify the expression of FP-tagged proteins. However, it comes with drawbacks, as this is a time-consuming experiment with a substantial cost due to the reagents required, and is not optimal for quantifications compared to fluorescence-based approaches. Here, we propose IGF detection as a fast, easy and efficient alternative to this technique.

Our study offers a comprehensive examination of IGF detection as an innovative tool for protein detection after SDS-PAGE. We provide a systematic comparison of a number of widely used FPs. Our evaluation considered key parameters such as fluorescence intensity, heat-stability, and their overall suitability for IGF detection in the context of standard SDS-PAGE protocols and accessible imaging devices. Importantly, this technique can be performed without major adaptation of classical protocols except that protein denaturation should be carried out at low temperatures.

Overall, IGF detection is also a cost-effective method. This technique eliminates the need for specific antibodies, which may even not be available in the case of less conventional FPs, as well as nitrocellulose membrane or ECL reagents. This makes it an attractive option for labs that operate under budget constraints. Because the observation is done directly from the gel after migration with no additional steps, problems frequently observed with immunoblotting can be avoided, such as heterogeneity of protein transfer, uneven antibody deposition, stains, and uneven ECL spreading on the membrane.

Moreover, IGF is a time-saving technique, bypassing the need for protein transfer and blotting/washing steps, as the proteins of interest can be visualized immediately after migration. For example, we were able to use this technique to quickly screen for GFP-containing clones directly on lysates from colonies after transformation (See Material and Methods). This complementary approach to PCR-based screening allows to quickly monitor protein expression and size in multiple clones. The advantages of IGF detection are even more appealing in the case of co-immunoprecipitations of two fluorescent proteins, since both the prey and the bait can be visualized within minutes after migration, without interference of primary antibody/secondary antibody recognition which can sometimes hinder signal analysis and quantification.

Normalization of the obtained IGF signal with respect to total proteins can be achieved without transfer and immunoblotting of loading controls by several means. This can involve commercial fluorescence-based approaches for total protein detection, after sample labeling (eg. Amersham QuickStain kit [Cytiva] or No-Stain™ protein labeling kit [Invitrogen]), or directly in gel (Stain-Free gels [Bio-Rad]). Also, after IGF imaging, a regular Coomassie staining of the gel can be reliably quantified in the near-infrared wavelengths (**Figure 9B**) (Butt and Coorssen, 2013).

Another key advantage of IGF detection is its versatility. All fluorescent proteins tested gave a fluorescent signal in gels, which is advantageous notably in the case of FPs for which antibodies are not yet available. However, not all FPs perform equally under all conditions. The results show variations in the efficiency of IGF detection, emphasizing the need to consider both the FP’s fluorescence properties and its resistance to denaturation. Our findings demonstrate that EGFP and sfGFP, in particular, display strong in-gel signals and good resistance to denaturation. These features make them prime candidates for IGF applications. Moreover, mCherry and mKO-κ stand out for their exceptional heat-stability, maintaining their fluorescence even at relatively high denaturing temperatures. This expands the possibilities of IGF detection to proteins that require higher denaturation temperatures (such as proteins forming dimers or assembled in tight complexes). In all cases, though, the intensity of the signal and the ability to specifically detect a fluorophore depend on the available filters on the imaging systems, which were sometimes limiting (especially in the case of mCherry) and could lead to bleed-through of signals when visualizing green- and red-tagged proteins in the same sample.

Since fluorescent signals originate from the protein itself, the IGF detection method reduces potential biases that can be encountered after using primary and secondary antibodies. Like all fluorescence detection protocols, signal detection does not depend on a chemiluminescence enzymatic reaction, which imposes kinetic constraints and whose activity can vary depending on substrate accessibility. Depending on the antibody used, and thanks to the constant improvement in the sensitivity of imaging devices, we found that IGF detection could outperform antibody-based chemiluminescent detection, and the signal was globally more linear with respect to protein concentration. This technique also allowed the detection of endogenous proteins with low abundance. Moreover, changing acquisition settings may increase sensitivity, with less background than observed with chemiluminescence.

However, like all techniques, IGF has its limitations. First, this technique relies on a mild denaturation of proteins, which is not compatible with harsh treatments such as TCA precipitation, which is often used to prepare yeast lysates. Whereas native lysate preparation may be more complex than a simple TCA precipitation, it does not require more material and overall this protocol remains faster considering that signals can be monitored immediately after migration. Moreover, our protocol is compatible with regular lysate preparation from mammalian cells which is often made in native conditions. Second, denaturation is performed at lower temperatures than usually performed in classical protocols. We demonstrated that this was sufficient to separate proteins from higher-order complexes (proteasome, mitochondrial ribosomes, RNA polymerase I) suggesting no apparent drawbacks at this level. However, the tagged proteins themselves may also be partially resistant to denaturation at the low temperatures used for IGF, which would lead to more than one fluorescent bands. This should be examined when setting up IGF for the first time on a given protein. Moreover, depending on the FP used for tagging, proteins treated in mild denaturation conditions can migrate as 2 discrete bands corresponding to the fluorescent and denatured forms of the FP. The apparent molecular weight of FP-tagged proteins is usually lower than under classical denaturation conditions, with up to 15-kDa difference with the denatured form in the case of mRuby2 or mKO-κ. Determining which temperature is best suited for a given FP may require setting up the conditions on a protein for which an antibody is available, so as to evaluate the extent of denaturation. This is an important step as this could preclude its use for absolute quantifications. The use of the most denaturation-resistant fluorescent proteins, such as sfGFP, mCherry or mKO-κ would be advised for these applications. Although IGF was compatible with a range of lysis buffers, it is also possible that the composition of the lysis buffer in which proteins are resuspended may alter overall fluorescence, depending on the abilities of FPs to remain folded and fluorescent in these conditions. It should be noted that we did not systematically analyze whether the linker region between the protein of interest and the fluorescence protein influenced IGF signal and FP denaturation. Finally, antibodies may recognize denatured and non-denatured forms of the FPs with different affinities. While these factors do not diminish the value of IGF, they highlight the need for appropriate controls when setting up IGF for protein detection.

The method’s simplicity, cost-effectiveness, reliability and compatibility with existing FPs and current protocols (including co-immunoprecipitation and SNAP-tagging) make IGF detection an attractive alternative to immunoblotting which can greatly benefit research in a range of fields. From basic molecular biology to more applied biomedical research, IGF offers a robust and economical solution to detect fluorescent proteins in gels. Future studies could further explore the method’s robustness across different cell types and conditions, ensuring its broad applicability in diverse experimental settings.

## Material and Methods

### Yeast strains and cultures

Strains are derivatives of the BY4741 background and are listed in Supplementary Table 1. Proteins were tagged with various FPs at their endogenous loci by homologous recombination using plasmids listed in Supplementary Table 2, or originated from the collection of GFP-tagged strains (Huh et al., 2003). All yeast strains were constructed by transformation with the standard lithium acetate–polyethylene glycol protocol using homologous recombination and verified by polymerase chain reaction (PCR) on genomic DNA. Yeast cells were grown in YPD medium (2% w/v) or in SC medium [containing yeast nitrogen base (1.7 g/L; MP Biomedicals), ammonium sulfate (5 g/liter, Sigma-Aldrich), the appropriate drop-out amino acid preparations (MP Biomedicals), and 2% (w/v) glucose, unless otherwise indicated]. Precultures were incubated at 30°C for 8 hours and diluted in the evening to 20-mL cultures to reach mid-log phase the next morning.

### Preparation of cell lysates from yeast

Exponentially growing cells (5 OD_600_ equivalents) were washed twice in water and resuspended in 100µL of cold native lysis buffer [Tris-HCl pH 8.0, 100 mM, NaCl, 0.15 M, Glycerol 5% v/v, Triton X-100 0.5% v/v, PMSF 0.5 mM, Complete antiprotease EDTA free (Roche, #11836170001)], and incubated for 5 min on ice-cold water with glass beads (Sigma-Aldrich, #G8772) to cool samples down prior to lysis. Cells were then lysed in a cold room on a vortex (4 x 30 sec, with 1 min incubation on ice-cold water between each lysis). Cell lysates were retrieved by piercing the bottom of the 1.5-mL tubes and brief centrifugation in a minispin centrifuge. Lysates were cleared by centrifugation at 4°C for 5 min at 3,000 xg. Supernatants were collected and 4X Laemmli sample buffer (Bio-Rad, #161-0747) containing DTT (12.5 µM final concentration) was added to the lysates (1X final concentration). Samples were denatured as indicated for each experiment, and loaded on a SDS-PAGE gel.

To use IGF from colonies growing on plates for screening purposes, 3 days-old colonies were harvested and resuspended in 100 µL cold native lysis buffer [Tris-HCl pH 8.0, 100 mM, NaCl, 0.15 M, Glycerol 5% v/v, Triton X-100 0.5% v/v, PMSF 0.5 mM, Complete antiprotease EDTA free (Roche, #11836170001)] and lysed as described above for exponentially growing cells. 15 µL of lysate were collected and 4X Laemmli sample buffer (Bio-Rad, #161-0747) containing DTT (12.5 µM final concentration) was added to the lysates (1X final concentration). Samples were denatured at 30°C for 5 min, and 20 µL were loaded on SDS-PAGE gel.

Relative quantitations were carried out with a detergent-compatible Bradford reagent (Abcam, #ab119216), and samples were then incubated for 5 min. Denaturation temperatures varied from 30°C to 95°C and are indicated in the Results section and the Figures. Equivalent protein amounts were loaded on commercial precast 4-20% TGX gels (Bio-Rad, #4561094), and run in Tris-glycine-SDS (TGS) buffer at 150 V for 45 min unless otherwise indicated.

### Preparation of yeast protein extracts with TCA

A volume of culture corresponding to 1 OD_600_ equivalent of exponentially growing yeast cells was collected, to which TCA (100%, w/v; Sigma-Aldrich) was added (10% final concentration). Whole cells were precipitated on ice for 10 min, samples were then centrifuged at 16,000 xg at 4°C for 10 min, the pellet was resuspended in 100 µL of a 10% (w/v) TCA solution and broken for 10 min with glass beads (Sigma-Aldrich, #G8772) at room temperature. Lysates were transferred to another 1.5-mL tube to remove glass beads and centrifuged for 5 min at 16,000 xg at 4°C. Protein pellets were resuspended in 50 µL of 1X Laemmli sample buffer, which included 50 mM Tris-base (final concentration) to buffer for the TCA present in the pellet.

### Preparation of cell lysates from Drosophila cultured cells

Drosophila wing imaginal disc cultured cells [Clone 8 (Cl-8) cells] responsive to Hedgehog (HH) were cultured and transiently transfected using transitory insect transfection reagent (Mirusbio, # MIR 6100) using pAct-EGFP, pAct-SMO-EGFP, and pAct-HH (Malpel et al., 2007; Sanial et al., 2017) as described previously (Sanial et al., 2017). Forty-eight hours after transfection, cells were washed in PBS and lysed in 1% Triton X-100, 150 mM NaCl, 50 mM Tris-HCl pH 8.0, 1 mM dithiothreitol with complete EDTA-free antiprotease mix (Roche, #11836170001) and PhosSTOP (Roche, #04906837001). Lysates were centrifuged (12,000 xg) for 10 min at 4°C and the supernatant was mixed with 4X Laemmli sample buffer (Bio-Rad) containing DTT (12.5 µM final concentration). Samples were incubated for 5 minutes at 25°C before loading on a homemade 10% acrylamide gel without SDS. Gels were run for 90 min at 150 V in TGS buffer on a MiniProtean setup (Bio-Rad).

### Preparation of cell lysates from fly ovary tissue

Ovary extracts were obtained from Ptub64c-GAL4 and Ptub64c-GAL4; UbiEGFP-PAR3 transgenic flies (Kullmann and Krahn, 2018) by dissecting ovaries (20 flies per genotype) into 1X PBS. Ovaries were placed on ice in lysis buffer [10 mM Tris-HCl pH 7.5, 150 mM NaCl, Complete Protease Inhibitor cocktail (Roche)] and mechanically homogenized using micro pestles in matching tubes. Ovaries lysates were spun at 3000 xg for 20 min at 4°C to eliminate debris. Supernatants were incubated for 5 min at 72°C in LDS sample buffer with Sample Reducing Agent (ThermoFisher, #NP0007). Samples were loaded on a NuPage, 4–12% Bis-Tris gels (ThermoFisher, #NP0322) and run in 1X MOPS buffer (ThermoFisher, #NP0001) for 90 min at 150v.

### Preparation of cell lysates from MDCK cells

MDCK cells (ATCC CCL-34) were transfected with Lipofectamine 2000 (Thermofisher #11668027) as described (Walch et al., 2018). Arf1-EGFP and Arf1-mCherry plasmids were obtained by subcloning Arf1 (Wessels et al., 2006) into pEGFP-N3 (Clontech) and pmCherry-N1 (Clontech), respectively. Cells were lysed in 6-well plates at confluency with 200 µL of lysis buffer [Tris-HCl pH 8.0 50 mM, NaCl 150mM, EDTA 10 mM, Triton X-100 0.5% (v/v) with complete antiprotease mix (Roche #11836145001)]. After 20 min incubation on ice, cells were scraped from the plate, collected in tubes and vortexed 3 x 5 sec, with 30 sec-incubation on ice between each step. Lysates were cleared by centrifugation 30 min, 11000 xg at 4°C. After protein quantitation using a Bradford reagent (Bio-Rad #5000205), samples were mixed with Laemmli sample buffer (Bio-Rad #161-0747) (1X final concentration). 50 µg of proteins were loaded per well on a precast 4-20%TGX gel (Bio-Rad, #4561094), and run in TGS buffer at 200 V for 45 min.

### Preparation of cell lysates from *C. elegans*

Worms [*C. elegans* strain N2 (wildtype ancestral, Bristol) and JDU233: *ijmSi63 [pJD520; mosII_5’mex-5_GFP::tba-2; mCherry::his-11; cb-unc-119(+)] II; unc-119(ed3) III* (Lacroix et al., 2024)] were cultured on MGM++ plate and fed on OP50 bacterial strain at 23C. Worm lysate was obtained from an asynchronous population. A total volume equivalent of 50 µL worm pellet was washed twice in bacterial M9 minimal medium containing 0.05% (v/v) Tween-20, resuspended into 200 µL Lysis buffer [50mM Tris-HCl pH 8.0, 200 mM NaCl, 5% glycerol and Complete protease inhibitor cocktail (Roche #11836145001)] and sonicated for 1 min on ice. Total extract was incubated for 5 min at 50°C with 1x final pH8 Laemmli sample buffer containing DTT (12.5 µM final concentration). 40 µg of worm extract were loaded per well onto a precast 4-20%TGX gel (Bio-Rad, #4561094), and run in TGS buffer at 200 V for 45 min.

### In-gel fluorescence imaging

After migration, the gel was imaged with gel-imaging systems whose excitation/emission wavelengths were compatible with the observation of green and red fluorescent proteins. Imaging with the Amersham Typhoon 5 (Cytiva) was carried out using the 488 nm laser with the Cy2 filter 525BP20 (for green FPs) or the 532 nm laser with the Cy3 filter LPG550 (for red FPs). The scanning resolution was 25 µm/pixel. PMT voltage was adjusted so that collected signals do not saturate. Imaging on the Chemidoc MP (Bio-Rad) was carried out using standard excitation/emission wavelengths recommended for Cy2/Cy3 filters (green FP: Epi-blue, excitation: 460−490 nm, emission: 518–546 nm; red FP: Epi-green, excitation: 520−545 nm, emission: 577–613 nm).

Total proteins were visualized by in-gel fluorescence using a trihalo compound incorporated in the commercial pre-cast SDS–PAGE gels (stain-free TGX gels, 4–20%; Bio-Rad) after 45 sec UV-induced photoactivation using a ChemiDoc MP imager (Bio-Rad), serving as a loading control. Note that UV irradiation should preferentially be performed after FP imaging (see **Figure S2**).

### Immunoblotting

After fluorescence imaging, gels were equilibrated in transfer buffer [25 mM Tris, 190 mM glycine, 20% ethanol (v/v), 0.02% SDS (w/v)] for 15 min and transferred at 100 V onto a nitrocellulose membrane in a liquid transfer system (MiniProtean, Bio-Rad). Membranes were blocked in Tris-buffered saline solution (50 mM Tris pH 7.6, 150 mM NaCl) containing 0.1 % Tween-20 (v/v) (TBS-T) and 2% fat-free milk (w/v) for 30 min and incubated for at least 2 hours with the appropriate primary antibodies. Membranes were washed 3 × 10 min in TBS-T and incubated for at least two hours with the corresponding secondary antibody (coupled with horseradish peroxidase or Alexa680). Membranes were then washed again 3 × 10 min in TBS-T and incubated with ECL Select Western Blotting Detection Reagent (Cytiva #RPN2235) except in **Figure S3B** in which Clarity Western ECL Substrate (Bio-Rad #1705060) was used because of the strong signal obtained with the anti-GFP antibody 3H9. Luminescence signals were acquired using a ChemiDoc MP (Bio-Rad). For denaturation of proteins on the membrane (**Figure S3B**), the membrane treatment was adapted from Xu et al (2019) (Xu et al., 2019): after transfer, the membrane was washed with water, incubated with cold 50% methanol (v/v) on ice for 30 min. The membrane was then washed with water, quickly dried on lint-free wipes and incubated in a sandwich between two sheets of Whatman paper and two glass slides at 75°C in a preheated thermocycler for 30 min.

### Antibodies

The primary antibodies used are: anti-GFP (clones 7.1 and 13.1; Roche #11814460001; 1/5,000 dilution), Anti-GFP (Chromotek #3H9; 1/5,000), anti-GFP-DyLight800 (Rockland, #600-145-215; 1/2,000), anti-Bmh1 (kind gift of S. Lemmon, Univ. Miami, USA; 1/15,000) (Gelperin et al., 1995); anti-Hkx2 (Rockland, #100-4159; 1/3,000). Secondary antibodies are anti-Mouse IgG-HRP (Sigma-Aldrich, #A5278; 1/5000), anti-Rabbit IgG-HRP (Sigma-Aldrich, #A6154; 1/5,000), anti-Rat IgG-HRP (Jackson Immuno, #112-035-143; 1/5,000), and anti-Mouse IgG Alexa Fluor 680 (ThermoFisher, #21057; 1/5,000).

### Test of the linearity of the signal by sample dilution

Extraction was performed in native conditions using 15 OD_600_ equivalent of yeast lysed in 300 µL of cold native lysis buffer [100 mM Tris-HCl pH 8.0, NaCl 0.15 M, Glycerol 5% v/v, Triton X-100 0.5% v/v, PMSF 0.5 mM, Complete antiprotease EDTA free (Roche, #11836170001)]. 4X Laemmli sample buffer (Biorad) was added to the lysates (1X final, 12.5 µM DTT final concentration), and samples were then incubated for 5 min at 30°C. Samples were serial-diluted with 1X native sample buffer (62.5 mM Tris-HCl pH 6.8, 10% glycerol (v/v), 0.005% bromophenol blue (w/v)], as follows: 9 times at 0.75X dilution, followed by 5 times at 0.5X dilution (final dilution >300 fold). Fluorescence levels were quantified using ImageQuant TL v8.2 (Cytiva).

### Sample denaturation with a range of temperatures

Extraction was performed in native conditions using 15OD_600_ equivalent of yeast lysed in 300µL of cold native lysis buffer [100 mM Tris-HCl pH 8.0, NaCl 0.15 M, Glycerol 5% v/v, Triton X-100 0.5% v/v, PMSF 0.5 mM, Complete antiprotease EDTA free (Roche, #11836170001)]. 4X Laemmli sample buffer was added to the lysates (1X final, DTT 12.5 µM final concentration). Samples were distributed in 0.5 mL Eppendorf tube and kept on ice until heat treatment. Samples were heated at various temperatures in an Eppendorf Gradient Mastercycler for 5 min, and kept at 22°C after incubation. The equivalent of 0.5 OD_600_ was loaded in each well. Fluorescence levels were quantified using ImageQuant TL v8.2 (Cytiva).

### Co-immunoprecipitations of fluorescent proteins from yeast lysates

Exponentially growing cells (40 OD_600_ equivalents) were washed twice in water and resuspended in 400 µL of cold IP lysis buffer (100 mM Tris-HCl pH8, NaCl 200 mM, Glycerol 5%, Triton X-100 0.1%, EDTA 1 mM) containing protease inhibitors [PMSF 0.5 mM, Complete antiprotease EDTA free (Roche, #11836170001), NaF 10 mM], and incubated for 5 min on ice-cold water with glass beads. Cells were then lysed in a cold room on a vortex (4 x 30 sec, with 1 min incubation on ice-cold water between each lysis). Relative quantitation was made with Bradford reagent to use equivalent amounts of proteins for co-immunoprecipitation. Samples were diluted 4:10 as per the manufacturer’s instructions with dilution buffer (10 mM Tris-HCl pH 8.0, NaCl 200 mM, EDTA 1 mM) containing protease inhibitors [PMSF 0.5 mM, Complete antiprotease EDTA free (Roche, #11836170001), NaF 10 mM]. A fraction (3%) was taken as the “Input” fraction. 10 µL GFP-Trap Magnetic Particles M-270 (Proteintech, #gtd) or RFP-Trap Magnetic Agarose (Proteintech, #rtma) were added to the remaining of the sample and incubated on a rotating wheel for 1h at 4°C and washed 3 times with cold IP lysis buffer. Proteins were eluted in 40 µL of FP trap sample buffer [66 mM Tris-HCl pH 8.0, 2% SDS, 100 mM NaCl, 12.5 µM DTT, 10% (v/v) glycerol, 0.002% bromophenol blue (w/v)] by incubating at 50°C for 20 min with soft shaking in a dry bath incubator. Half of the IP and 1% of the diluted lysate were loaded on the gel.

### SNAP-tagging

Yeast total lysates were prepared as in “Preparation of cell lysates from yeast” using cold native lysis buffer supplemented with DTT (0.5 mM final concentration). 20 µL (corresponding to 1 OD equivalent) of extracts were incubated for 30 min at 37°C with 10 µL of staining solution [Tris-HCl pH 8.0 100 mM, DTT 0.5 mM, PMSF 0.5 mM, Complete antiprotease EDTA free (Roche, #11836170001)] in the presence or absence of SNAP-surface Alexa fluor 647 subtrate (New England Biolabs, # S9136S) at a final concentration of 1.66 µM. LDS sample buffer was added (1X final) and samples were denatured for 5 min at 30°C before loading onto a commercial pre-cast SDS–PAGE gels (stain-free TGX gels, 4–20%; Bio-Rad).

### Construction of yeast-optimized mCherry, mNeonGreen and FP-tagging vectors

The mCherry and mNeonGreen sequences were codon-optimized for *Saccharomyces cerevisiae* using GENEius tool (giving ymCherry and ymNeonGreen, respectively) and synthetized (Eurofins Genomics), with a GSAGAGAGAGAG or GSGAGAGAGAGAGA linker respectively, flanked by the traditional S1/S3 oligo sequences used for amplification (Janke et al., 2004). Linkers were also codon-optimized but sequences are different to avoid recombination. DNA was provided cloned into pEX-A128 containing a polylinker site to allow subcloning into the pYM vector series (Janke et al., 2004) with various selection markers. ymNeonGreen was cloned in pYM16 at BsiW1/AscI sites, giving pYM-ymCherry-NAT (pSL778), and ymCherry was cloned in pYM17 at HindIII/AscI sites, giving pYM-ymNeonGreen-Hph (pSL766). Tagging with sfGFP was achieved using the GTH-g plasmid (gift from Serge Pelet; Addgene #81104) (Wosika et al., 2016). Tagging with mNeonGreen was achieved using pFA6-mNeongreen (HPH) plasmid (kind gift from Silke Hauf). Tagging with mKO-κ was achieved using pYM GAGAGA ymKOKappa-hygro plasmid (pMC215, kind gift from Aurélie Massoni). Tagging with yeast codon-optimized Tag-RFP-T or mRuby2 was achieved using the pFa6-link-yoTag-RFP-T-CaURA3 and pFA6a-link-yomRuby2-SpHIS5, respectively (gifts from <u>Kurt Thorn</u> and <u>Wendell Lim</u>, Addgene plasmids #44877 and #44843). Tagging with EGFP was made using pYM27 (Janke et al., 2004) or by switching the GFP(S65T)-HIS3MX cassette from strains originating from the GFP collection (Huh et al., 2003) to an EGFP-KanMX cassette by homologous recombination. Tagging with SNAP was achieved using pBS-SKII-3XHA-fSNAP-NAT (gift from Stephen Buratowski, Addgene plasmid #188916) (Baek et al., 2022).

### Correlation of published protein abundance with IGF signals

Data of protein abundance were published in Ho et al., 2018 (Ho et al., 2018) and retrieved from the Saccharomyces Genome Database (www.yeastgenome.org). For Figure 7C, average abundances (±SD) were calculated for the indicated proteins using data available from non-treated/unchallenged cells, and plotted against the signal intensity obtained by IGF measurement normalized to total protein staining (n=3). For Figure S8, data were retrieved for abundance studies based on mass spectrometry, confocal microscopy or immunoblotting and averages were calculated (±SD when n≥3). Peason’s correlation test was calculated for each pair (abundance and IGF signal) using GraphPad Prism.

## Acknowledgments

The authors wish to thank Dr Alenka Copic (CRBM, Montpellier, FR), Dr Julien Dumont (Institut Jacques Monod, Paris, FR), Dr Silke Hauf (Virginia Tech, Blacksburg, VA, USA), Dr Sandra Lemmon (University of Miami, Coral Gables, FL, USA), Mrs Aurélie Massoni (IBGC, Bordeaux, France) and Dr Myriam Ruault (Institut Curie, Paris, France) for sharing material, and Dr Catherine L. Jackson (Institut Jacques Monod, Paris, FR) for discussions and critical reading of the manuscript. This work was supported by grants from Université Paris-Cité IDEx (https://u-paris.fr; ANR-18-IDEX-0001, “Emergence en Recherche” RS30J20IDXA1_EMERLEON) and Agence Nationale pour la Recherche (“AMPKILL”, ANR-23-CE13-0012-01) to S. Léon. N. Joly is supported by a grant from the Fondation ARC pour la recherche sur le cancer (ARCPJA2022050005002) and a grant from Université Paris-Cité IDEx (https://u-paris.fr; ANR-18-IDEX-0001, “Emergence en Recherche” RS30J23IDX64_KATAREP).

The authors declare no competing interests.

## Supplementary material

### Supplementary tables

**Table S1:**
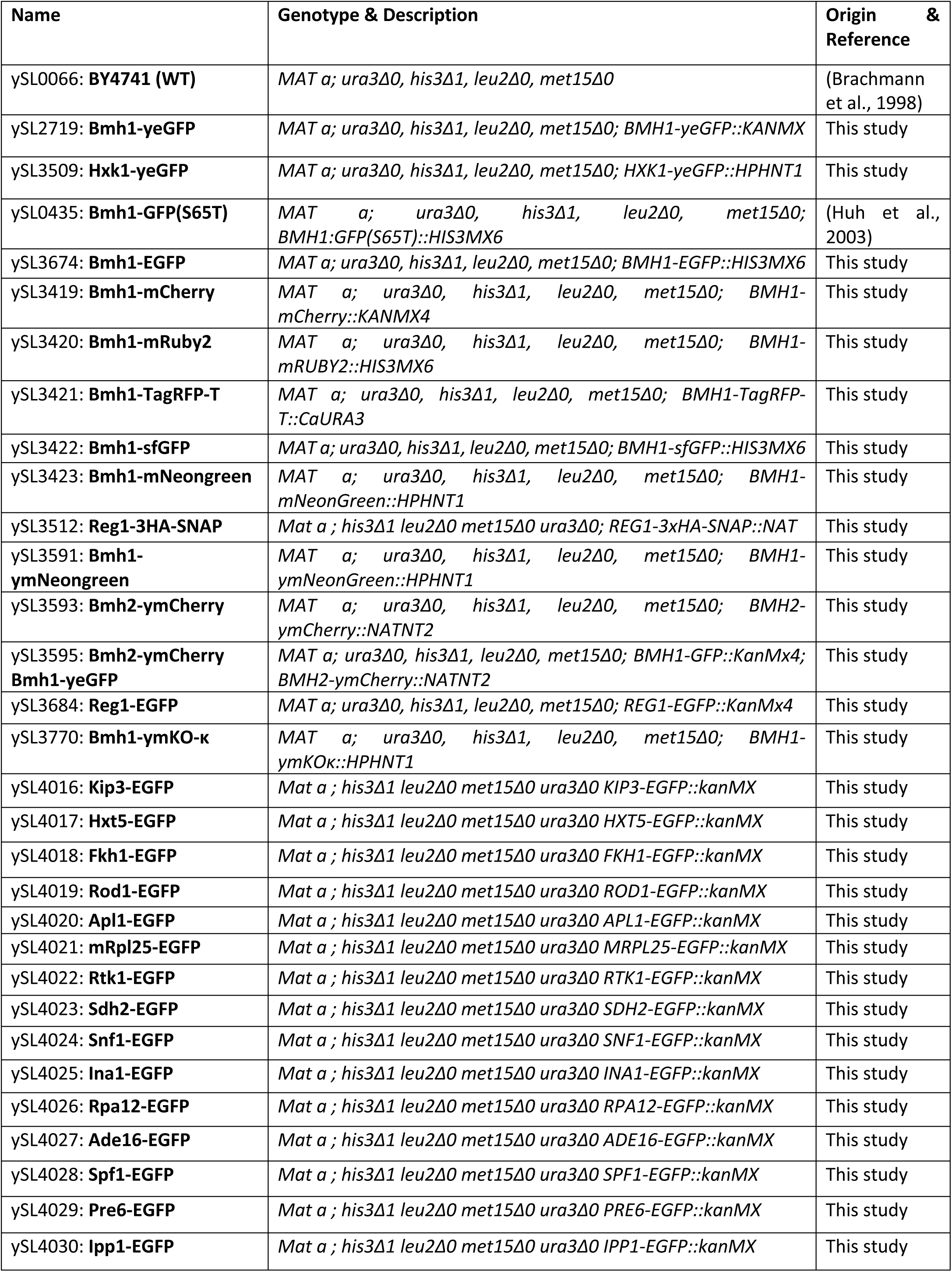
Yeast strains used in this study.

| Name | Genotype & Description | Origin & Reference |
| --- | --- | --- |
| ySL0066: **BY4741 (WT)** | *MAT a; ura3Δ0, his3Δ1, leu2Δ0, met15Δ0* | (Brachmann et al., 1998) |
| ySL2719: **Bmh1-yeGFP** | *MAT a; ura3Δ0, his3Δ1, leu2Δ0, met15Δ0; BMH1-yeGFP::KANMX* | This study |
| ySL3509: **Hxk1-yeGFP** | *MAT a; ura3Δ0, his3Δ1, leu2Δ0, met15Δ0; HXK1-yeGFP::HPHNT1* | This study |
| ySL0435: **Bmh1-GFP(S65T)** | *MAT a; ura3Δ0, his3Δ1, leu2Δ0, met15Δ0; BMH1:GFP(S65T)::HIS3MX6* | (Huh et al., 2003) |
| ySL3674: **Bmh1-EGFP** | *MAT a; ura3Δ0, his3Δ1, leu2Δ0, met15Δ0; BMH1-EGFP::HIS3MX6* | This study |
| ySL3419: **Bmh1-mCherry** | *MAT a; ura3Δ0, his3Δ1, leu2Δ0, met15Δ0; BMH1-mCherry::KANMX4* | This study |
| ySL3420: **Bmh1-mRuby2** | *MAT a; ura3Δ0, his3Δ1, leu2Δ0, met15Δ0; BMH1-mRUBY2::HIS3MX6* | This study |
| ySL3421: **Bmh1-TagRFP-T** | *MAT a; ura3Δ0, his3Δ1, leu2Δ0, met15Δ0; BMH1-TagRFP-T::CaURA3* | This study |
| ySL3422: **Bmh1-sfGFP** | *MAT a; ura3Δ0, his3Δ1, leu2Δ0, met15Δ0; BMH1-sfGFP::HIS3MX6* | This study |
| ySL3423: **Bmh1-mNeongreen** | *MAT a; ura3Δ0, his3Δ1, leu2Δ0, met15Δ0; BMH1-mNeonGreen::HPHNT1* | This study |
| ySL3512: **Reg1-3HA-SNAP** | *Mat a ; his3Δ1 leu2Δ0 met15Δ0 ura3Δ0; REG1-3xHA-SNAP::NAT* | This study |
| ySL3591: **Bmh1-ymNeongreen** | *MAT a; ura3Δ0, his3Δ1, leu2Δ0, met15Δ0; BMH1-ymNeonGreen::HPHNT1* | This study |
| ySL3593: **Bmh2-ymCherry** | *MAT a; ura3Δ0, his3Δ1, leu2Δ0, met15Δ0; BMH2-ymCherry::NATNT2* | This study |
| ySL3595: **Bmh2-ymCherry Bmh1-yeGFP** | *MAT a; ura3Δ0, his3Δ1, leu2Δ0, met15Δ0; BMH1-GFP::KanMx4; BMH2-ymCherry::NATNT2* | This study |
| ySL3684: **Reg1-EGFP** | *MAT a; ura3Δ0, his3Δ1, leu2Δ0, met15Δ0; REG1-EGFP::KanMx4* | This study |
| ySL3770: **Bmh1-ymKO-κ** | *MAT a; ura3Δ0, his3Δ1, leu2Δ0, met15Δ0; BMH1-ymKOκ::HPHNT1* | This study |
| ySL4016: **Kip3-EGFP** | *Mat a ; his3Δ1 leu2Δ0 met15Δ0 ura3Δ0 KIP3-EGFP::kanMX* | This study |
| ySL4017: **Hxt5-EGFP** | *Mat a ; his3Δ1 leu2Δ0 met15Δ0 ura3Δ0 HXT5-EGFP::kanMX* | This study |
| ySL4018: **Fkh1-EGFP** | *Mat a ; his3Δ1 leu2Δ0 met15Δ0 ura3Δ0 FKH1-EGFP::kanMX* | This study |
| ySL4019: **Rod1-EGFP** | *Mat a ; his3Δ1 leu2Δ0 met15Δ0 ura3Δ0 ROD1-EGFP::kanMX* | This study |
| ySL4020: **Apl1-EGFP** | *Mat a ; his3Δ1 leu2Δ0 met15Δ0 ura3Δ0 APL1-EGFP::kanMX* | This study |
| ySL4021: **mRpl25-EGFP** | *Mat a ; his3Δ1 leu2Δ0 met15Δ0 ura3Δ0 MRPL25-EGFP::kanMX* | This study |
| ySL4022: **Rtk1-EGFP** | *Mat a ; his3Δ1 leu2Δ0 met15Δ0 ura3Δ0 RTK1-EGFP::kanMX* | This study |
| ySL4023: **Sdh2-EGFP** | *Mat a ; his3Δ1 leu2Δ0 met15Δ0 ura3Δ0 SDH2-EGFP::kanMX* | This study |
| ySL4024: **Snf1-EGFP** | *Mat a ; his3Δ1 leu2Δ0 met15Δ0 ura3Δ0 SNF1-EGFP::kanMX* | This study |
| ySL4025: **Ina1-EGFP** | *Mat a ; his3Δ1 leu2Δ0 met15Δ0 ura3Δ0 INA1-EGFP::kanMX* | This study |
| ySL4026: **Rpa12-EGFP** | *Mat a ; his3Δ1 leu2Δ0 met15Δ0 ura3Δ0 RPA12-EGFP::kanMX* | This study |
| ySL4027: **Ade16-EGFP** | *Mat a ; his3Δ1 leu2Δ0 met15Δ0 ura3Δ0 ADE16-EGFP::kanMX* | This study |
| ySL4028: **Spf1-EGFP** | *Mat a ; his3Δ1 leu2Δ0 met15Δ0 ura3Δ0 SPF1-EGFP::kanMX* | This study |
| ySL4029: **Pre6-EGFP** | *Mat a ; his3Δ1 leu2Δ0 met15Δ0 ura3Δ0 PRE6-EGFP::kanMX* | This study |
| ySL4030: **Ipp1-EGFP** | *Mat a ; his3Δ1 leu2Δ0 met15Δ0 ura3Δ0 IPP1-EGFP::kanMX* | This study |

**Table S2:** Plasmids for yeast tagging used in this study.

| Name | Description | Origin & Reference |
| --- | --- | --- |
| **pYM12** | Plasmid to tag genes with yeGFP at their endogenous locus (*KANMX4*) | (Janke et al., 2004) |
| **pYM25** | Plasmid to tag genes with EGFP at their endogenous locus (*HPHNT1*) | (Janke et al., 2004) |
| **pYM28** | Plasmid to tag genes with EGFP at their endogenous locus (*HIS3MX6*) | (Janke et al., 2004) |
| **GTH-g (pSL648)** | Plasmid to tag genes with sfGFP at their endogenous locus (*HIS3MX6*) | Addgene #81104 (Wosika et al., 2016) |
| **pFA6-mNeongreen::Hph (pSL643)** | Plasmid to tag genes with mNeonGreen at their endogenous locus (*HPHNT1*) | Silke Hauf's lab, Virginia Tech; unpublished |
| **pFA6-link-yoTag-RFP-T-CaURA3 (pSL558)** | Plasmid to tag genes with yeast codon-optimized Tag-RFP-T at their endogenous locus (*CaURA3*) | Addgene #44877 (Lee et al., 2013) |
| **pFA6a-link-yomRuby2-SpHIS5 (pS649)** | Plasmid to tag genes with yeast codon-optimized mRuby2 at their endogenous locus (*SpHIS5*) | Addgene #44843 (Lee et al., 2013) |
| **pYM-mCherry (pSL1)** | Plasmid to tag genes with mCherry at their endogenous locus (*KANMX4*) | (Leon et al., 2008) |
| **pYM-ymCherry-NAT (pSL778)** | Plasmid to tag genes with yeast codon-optimized mCherry (*NATNT2*) | This study |
| **pYM-ymNeonGreen-Hph (pSL766)** | Plasmid to tag genes with yeast codon-optimized mNeonGreen (*HPHNT1*) | This study |
| **pYM ymKO-κ-Hph (pMC215)** | Plasmid to tag genes with yeast codon-optimized mKO-κ (*HPHNT1*) | Aurélie Massoni, IBGC, France; unpublished |

### Supplementary Figure

**Supplementary Figure 1.**
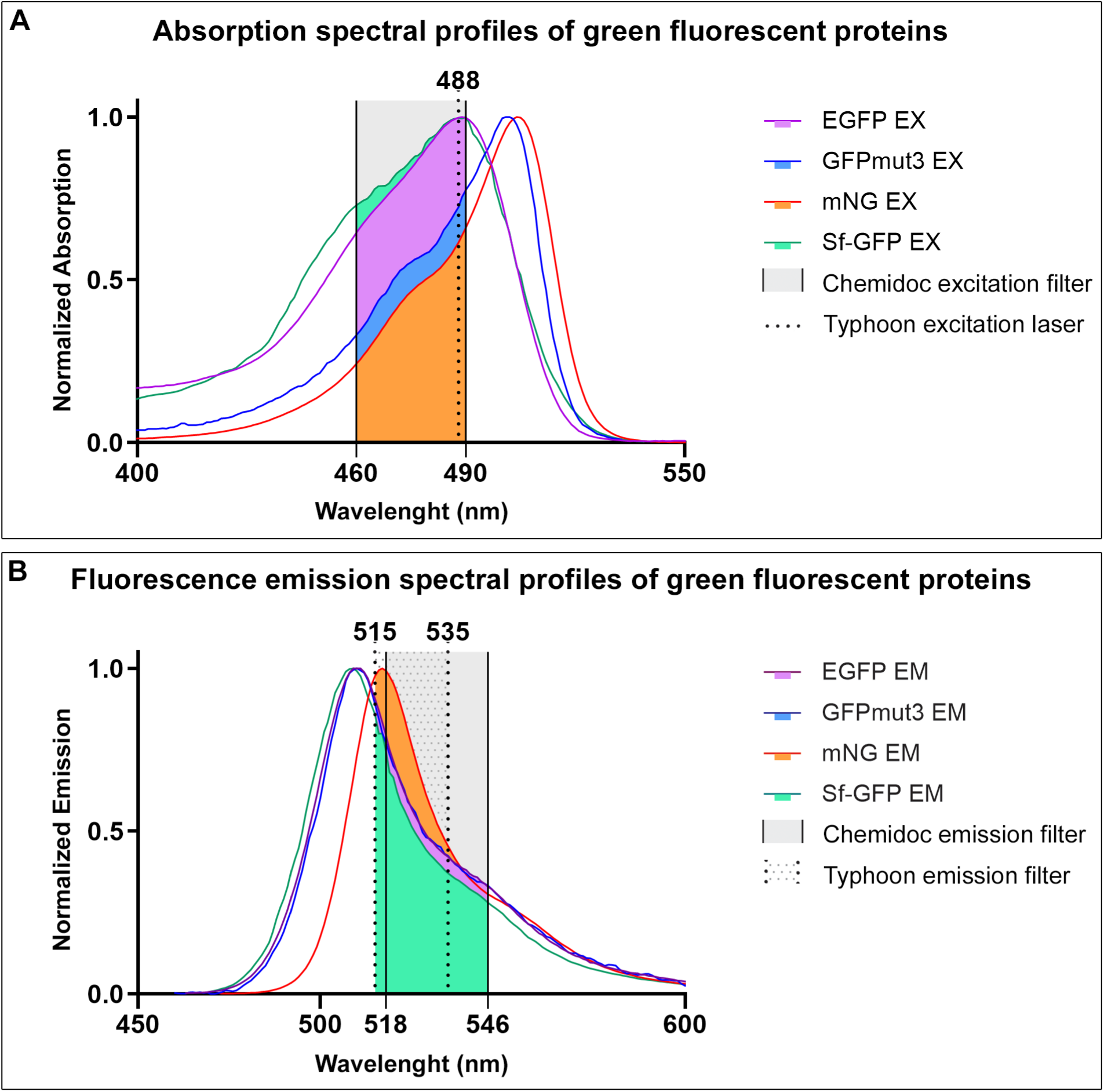
Spectral properties of green fluorescent proteins used in this study. A. Excitation wavelengths of yeGFP (= GFPmut3), EGFP, sfGFP and mNeonGreen. Data were retrieved from FPBase (Lambert, 2019). The window of excitation provided by the Chemidoc MP imaging system is indicated in gray (460-490nm), the wavelength of the excitation laser of the Typhoon is indicated as a dotted line (488 nm), as per the manufacturer’s indications. B. Emission wavelengths of yeGFP (=GFPmut3), EGFP, sfGFP, and mNeonGreen. Data were retrieved from FPBase (Lambert, 2019). The window of emission collected by the Chemidoc MP system is indicated in gray (518-546 nm), that of the Typhoon is indicated as shaded (515-535 nm), as per the manufacturer’s indications.

**Supplementary Figure 2.**
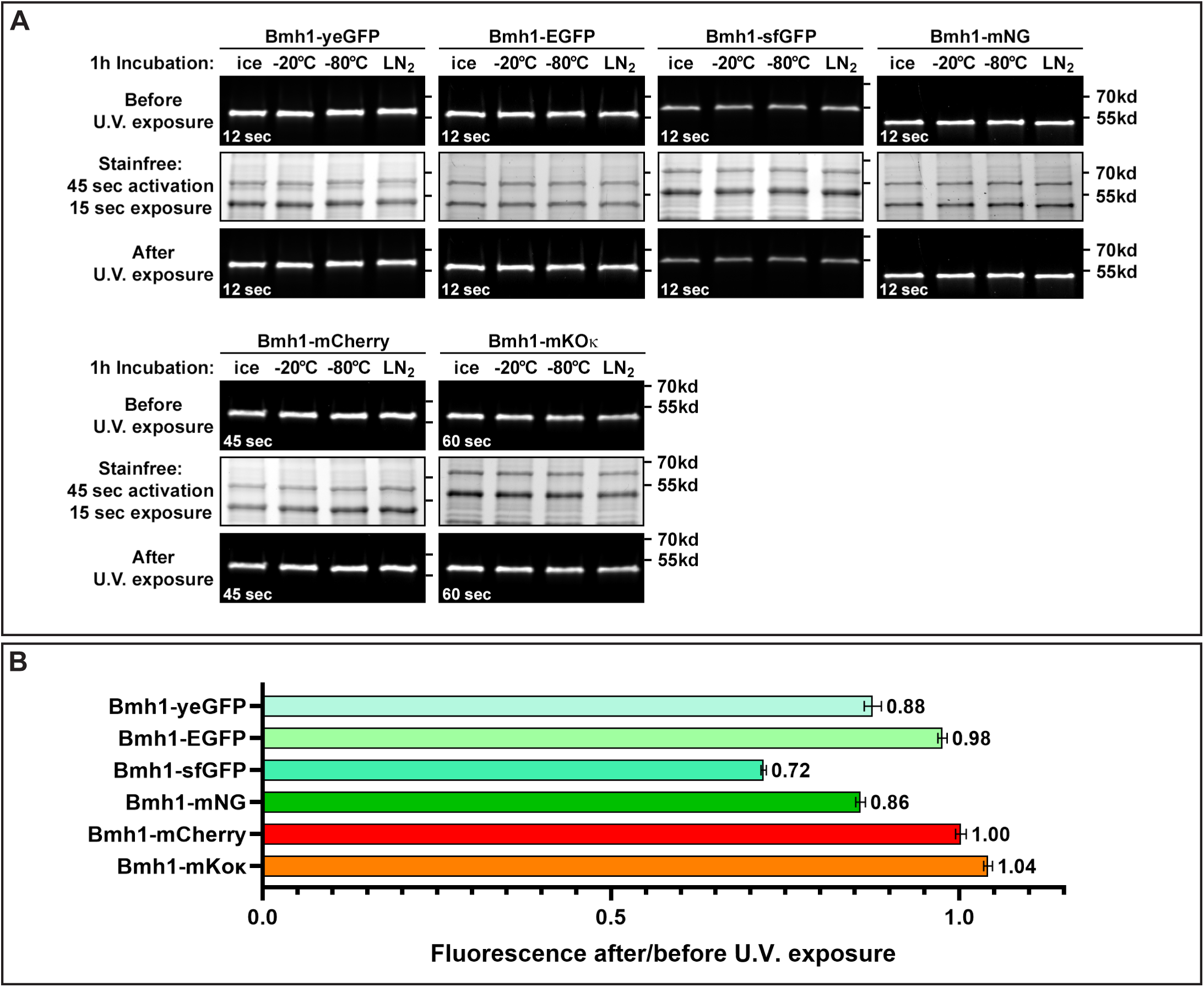
Effect of UV irradiation and freezing on the fluorescence of various fluorescent proteins. A. Native lysates of cells expressing Bmh1 tagged with yeGFP, EGFP, sfGFP, mNeonGreen, mCherry or mKO-κ were kept on ice or frozen (-20°C, -80°C, or in liquid N_2_) for one hour, thawed on ice and incubated at 30°C for 5 min, and loaded onto SDS-PAGE. IGF was detected on a Chemidoc MP before (top) and after (bottom) UV irradiation, which is required for the “StainFree” labeling (Bio-Rad) of total proteins with a trihalo compound (see Material and Methods). B. Quantification of the effect of UV exposure on IGF (ratio of signal obtained after UV exposure to that obtained before UV exposure, for each fluorophore; n=4; ± SD).

**Supplementary Figure 3.**
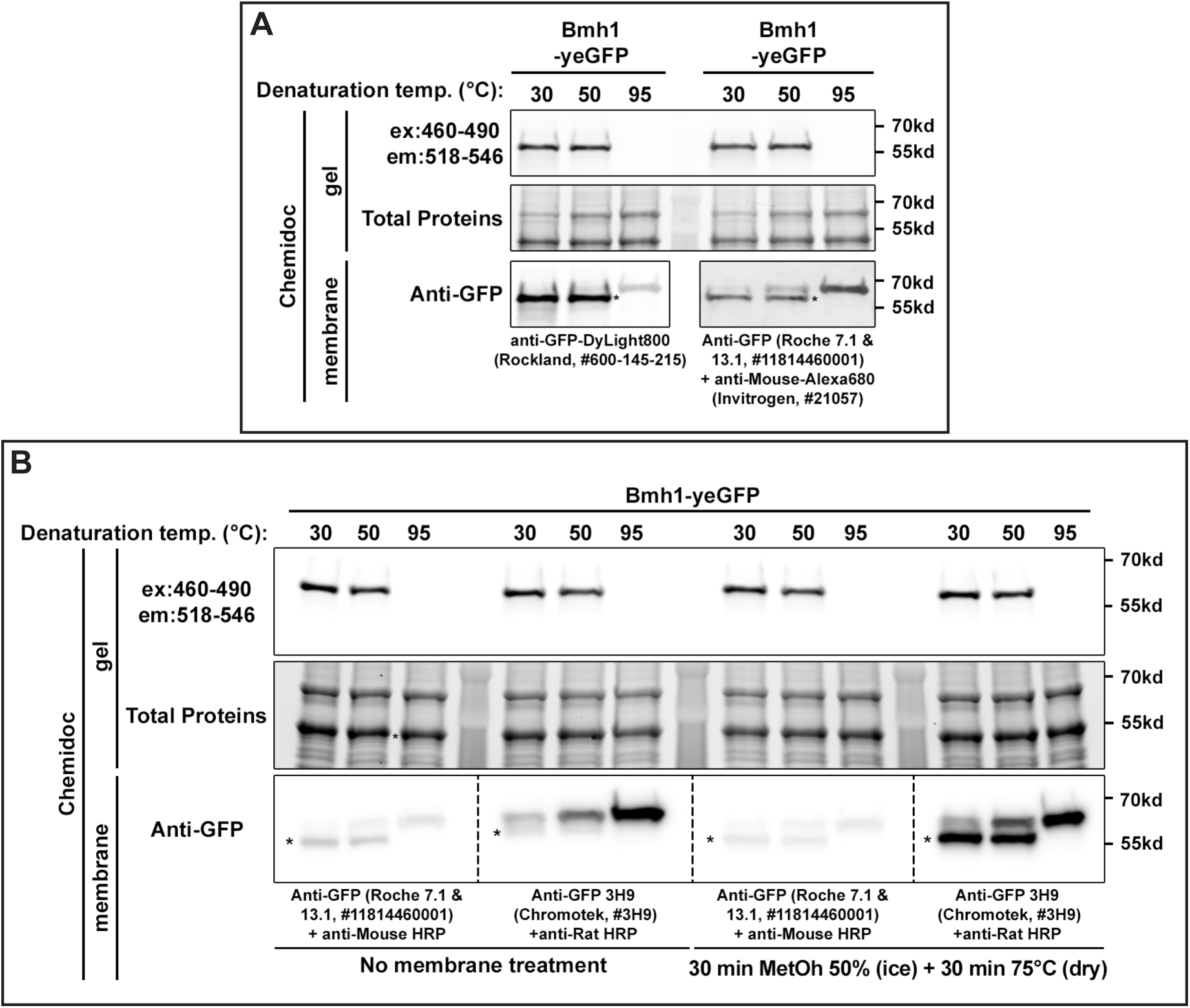
Effect of denaturation temperature on the recognition of GFP-tagged proteins by various anti-GFP antibodies. A. A native lysate of cells expressing Bmh1-yeGFP was resuspended in LDS sample buffer and incubated at the indicated temperatures. After migration, gels were imaged for green fluorescence using a Chemidoc MP, and total proteins were visualized by the stain-free technology on a Chemidoc MP. Proteins were then transferred to a nitrocellulose membrane, immunoblotted with the indicated antibodies and revealed using fluorescent secondary antibodies on a Chemidoc MP. * indicates the fluorescent species. B. A native lysate of cells expressing Bmh1-yeGFP was resuspended in LDS sample buffer and incubated at the indicated temperatures. After migration, gels were imaged for green fluorescence using a Chemidoc MP, and total proteins were visualized by the stain-free technology on a Chemidoc MP. Proteins were then transferred to a nitrocellulose membrane and immunoblotted with the indicated antibodies and revealed by chemiluminescence on a Chemidoc MP. Membranes on the right were treated with methanol and high temperature (see Material and Methods) to denature proteins and ameliorate the detection of GFP-tagged proteins treated at low temperatures. * indicates the fluorescent species.

**Supplementary Figure 4.**
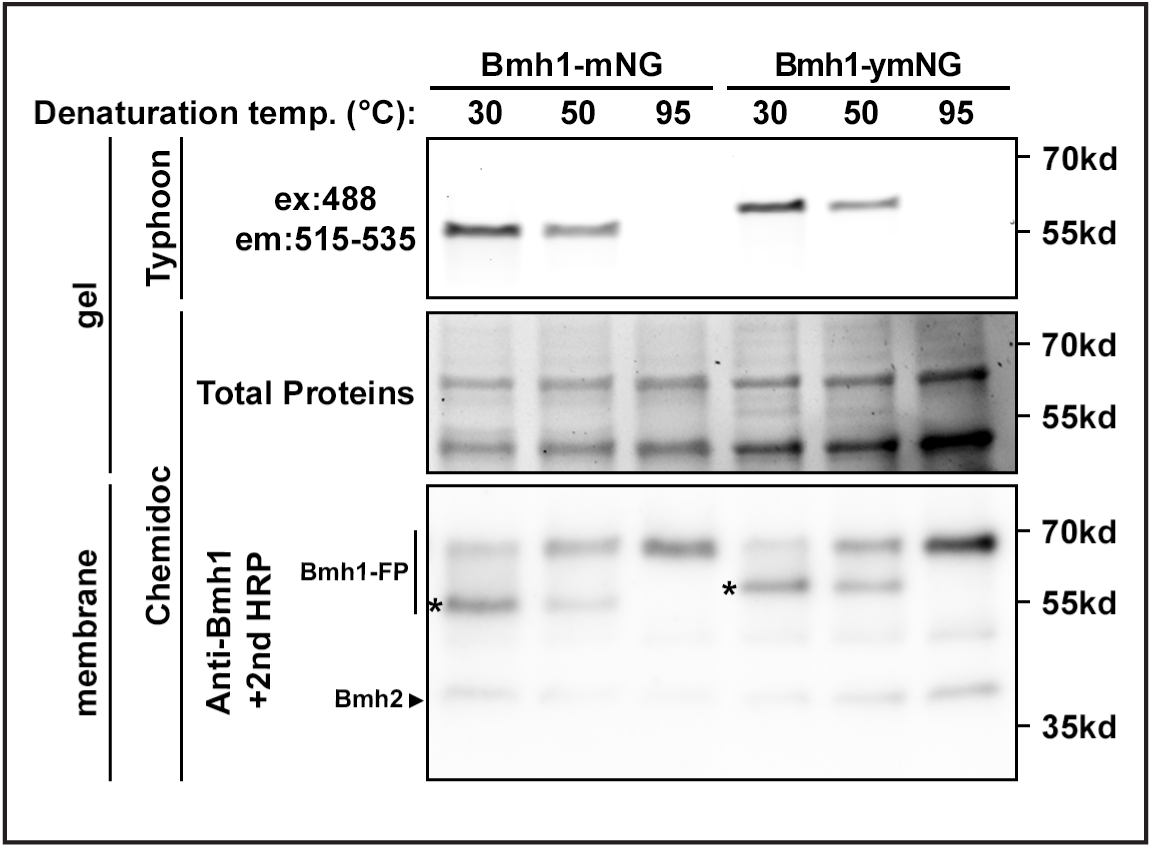
Comparison of IGF signals of mNeonGreen and a corresponding yeast-optimized version. Yeast expressing Bmh1-mNG and Bmh1-ymNG were lysed in native conditions, samples were resuspended in LDS sample buffer and incubated for 5 min at the indicated temperatures. After migration, gels were imaged for green fluorescence using a Typhoon or a Chemidoc MP, and total proteins were visualized by the stain-free technology on a Chemidoc MP. Proteins were then transferred to a nitrocellulose membrane and immunoblotted with anti-Bmh1 antibodies and then with anti-rabbit antibodies coupled to HRP, and revealed by chemiluminescence on a Chemidoc MP. The size difference is due to a longer linker present in the ymNeonGreen construct.* indicates the fluorescent species.

**Supplementary Figure 5.**
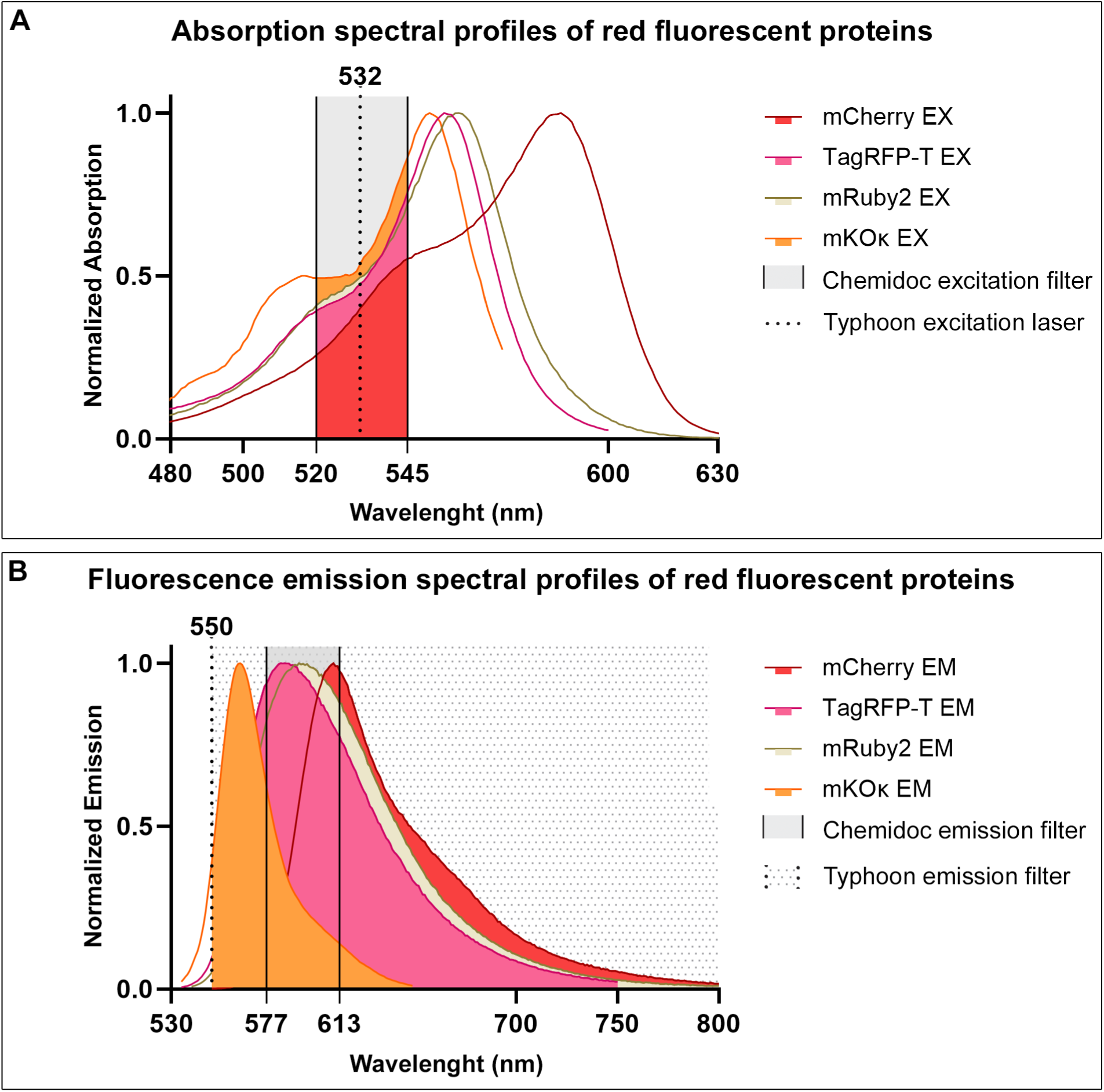
Spectral properties of red and orange fluorescent proteins used in this study. A. Excitation wavelengths of mCherry, TagRFP-T, mRuby2 and mKO-κ. Data were retrieved from FPBase (Lambert, 2019). The window of excitation provided by the Chemidoc MP imaging system is indicated in gray (520-545nm), the wavelength of the excitation laser of the Typhoon is indicated as a dotted line (532 nm), as per the manufacturer’s indications. B. Emission wavelengths of mCherry, TagRFP-T, mRuby2 and mKO-κ. Data were retrieved from FPBase (Lambert, 2019). The window of emission collected by the Chemidoc MP system is indicated in gray (577-613nm), that of the Typhoon is indicated as shaded (≥550 nm), as per the manufacturer’s indications.

**Supplementary Figure 6.**
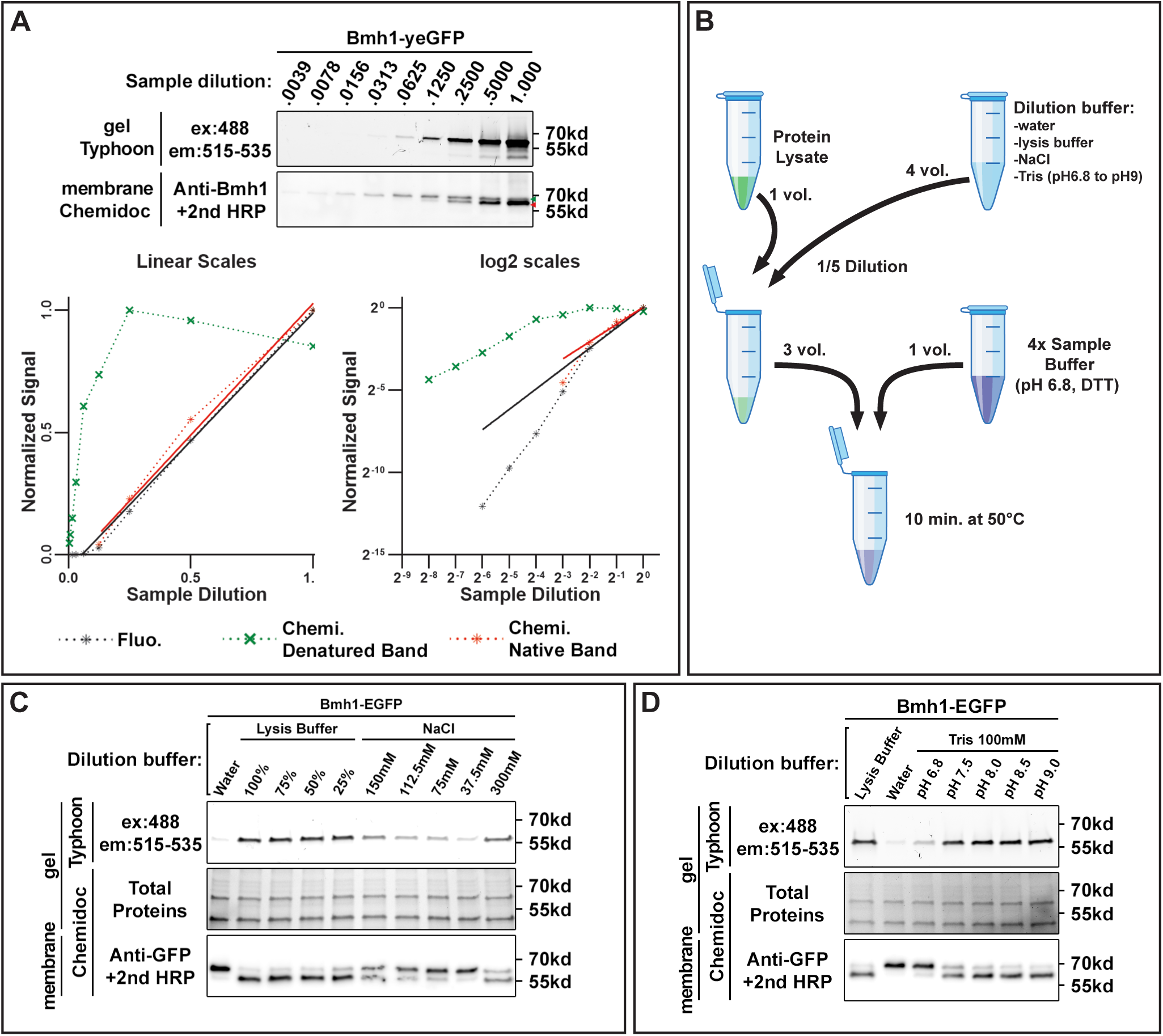
Effect of sample dilution on FP denaturation. A. Fluorescence of yeGFP is not maintained in LDS sample buffer at low protein concentrations. *Top*, Yeast expressing Bmh1-yeGFP were lysed in native conditions, samples were resuspended in LDS sample buffer, incubated for 5 min at 30°C and serially diluted (1:2) in LDS sample buffer. After migration, gels were imaged for green fluorescence using a Typhoon. Proteins were then transferred to a nitrocellulose membrane and immunoblotted with anti-Bmh1 antibodies and then with anti-rabbit antibodies coupled to HRP, and revealed by chemiluminescence on a Chemidoc MP. Red arrowhead: size of the native (fluorescent) protein; green arrowhead, size of the denatured protein. *Bottom*, Quantification of the signals obtained for fluorescence (black) and chemiluminescence (red: native protein, green: denatured protein) as a function of sample dilution. Linear-scaled and log-plot scaled graphs are shown. yeGFP denaturation increases with sample dilution, causing a bias when quantifying fluorescence. B. Design of the experiment to setup conditions to maintain FP fluorescence in diluted samples. Protein lysates prepared in native conditions from yeast expressing Bmh1-EGFP are diluted 1:5 with dilution buffer (water, lysis buffer, NaCl at various concentrations, or 100 mM Tris-HCl solution at various pH) before mixing with 4X sample buffer pH6.8 (1X final concentration). Samples are then incubated at 50°C for 10 min before being loaded on SDS-PAGE. C. Effect of lysis buffer and NaCl concentration on in-gel fluorescence. Samples were diluted with water, native lysis buffer (1X, 0.75X, 0.5X or 0.25X in water) or NaCl solution (at the indicated concentration) before being incubated at 50°C for 10 min. After migration on a commercial precast 4-20%TGX gel (Bio-Rad), gels were imaged for green fluorescence using a Typhoon, and total proteins were visualized by the stain-free technology on a Chemidoc MP. Proteins were then transferred to a nitrocellulose membrane and immunoblotted with anti-GFP antibodies and then with anti-mouse antibodies coupled to HRP, and revealed by chemiluminescence on a Chemidoc MP. D. Effect of pH on in-gel fluorescence. Samples were diluted with native lysis buffer (1X), water or Tris-HCl 100 mM solution (final concentration) at the indicated pH before being incubated at 50°C for 10 min. After migration on a commercial precast 4-20%TGX gel (Bio-Rad), gels were imaged for green fluorescence using a Typhoon, and total proteins were visualized by the stain-free technology on a Chemidoc MP.

**Supplementary Figure 7.**
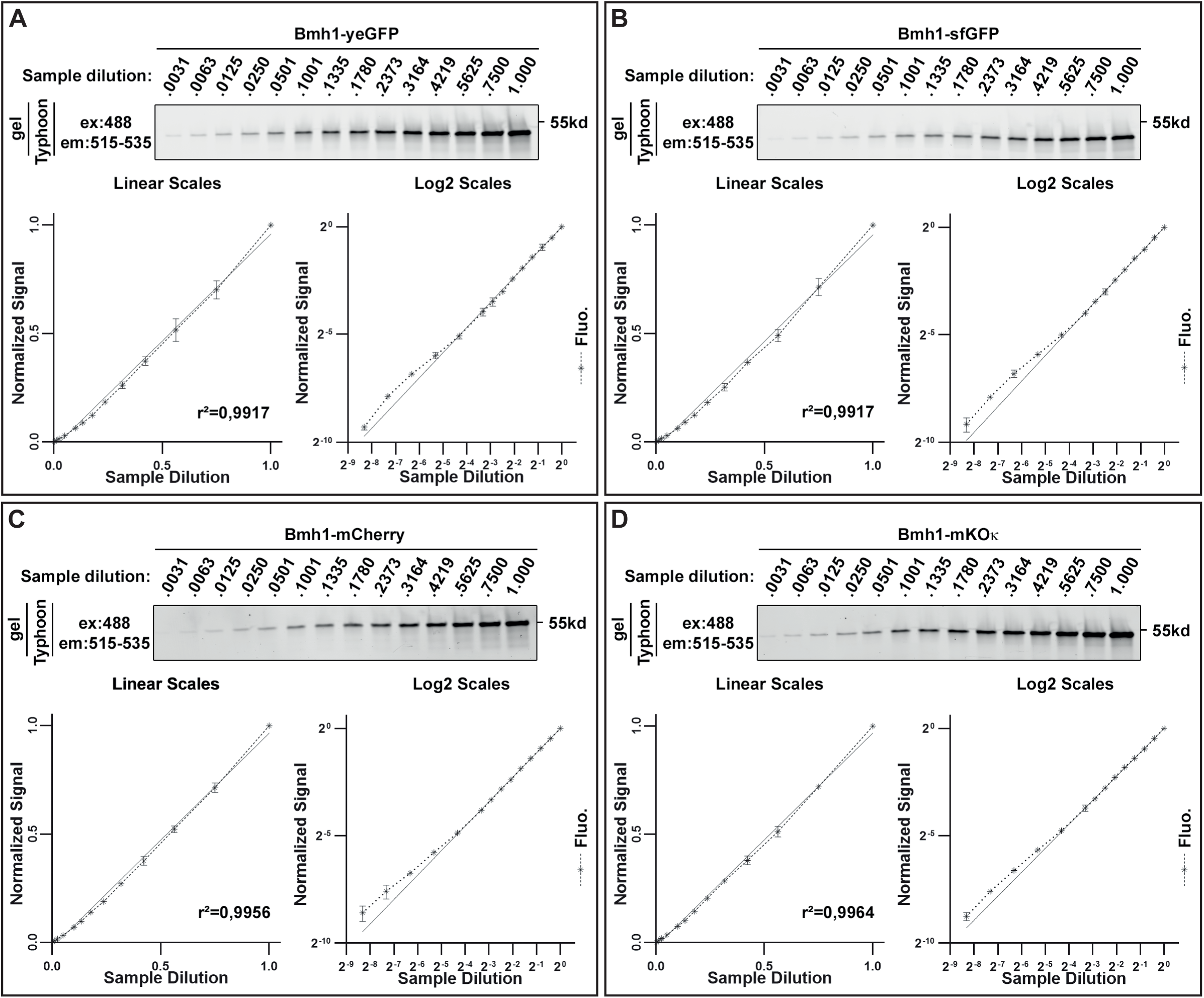
Sensitivity and linearity of in-gel fluorescence detection of various FP. A. Yeast expressing Bmh1-yeGFP were lysed in native conditions, samples were resuspended in LDS sample buffer and incubated for 5 min at 30°C. Samples were serially diluted (right to left) into sample buffer without LDS (see Material and Methods), and loaded onto SDS-PAGE. After migration, gels were imaged for fluorescence using a Typhoon. Quantification of the signals obtained for green fluorescence (black) and chemiluminescence (red) as a function of sample dilution (n=3 ± SD). Linear-scaled and log-plot scaled graphs are shown. B. Same as A, with Bmh1-sfGFP. C. Same as A, with Bmh1-mCherry. D. Same as A, with Bmh1-mKO-κ.

**Supplementary Figure 8.**
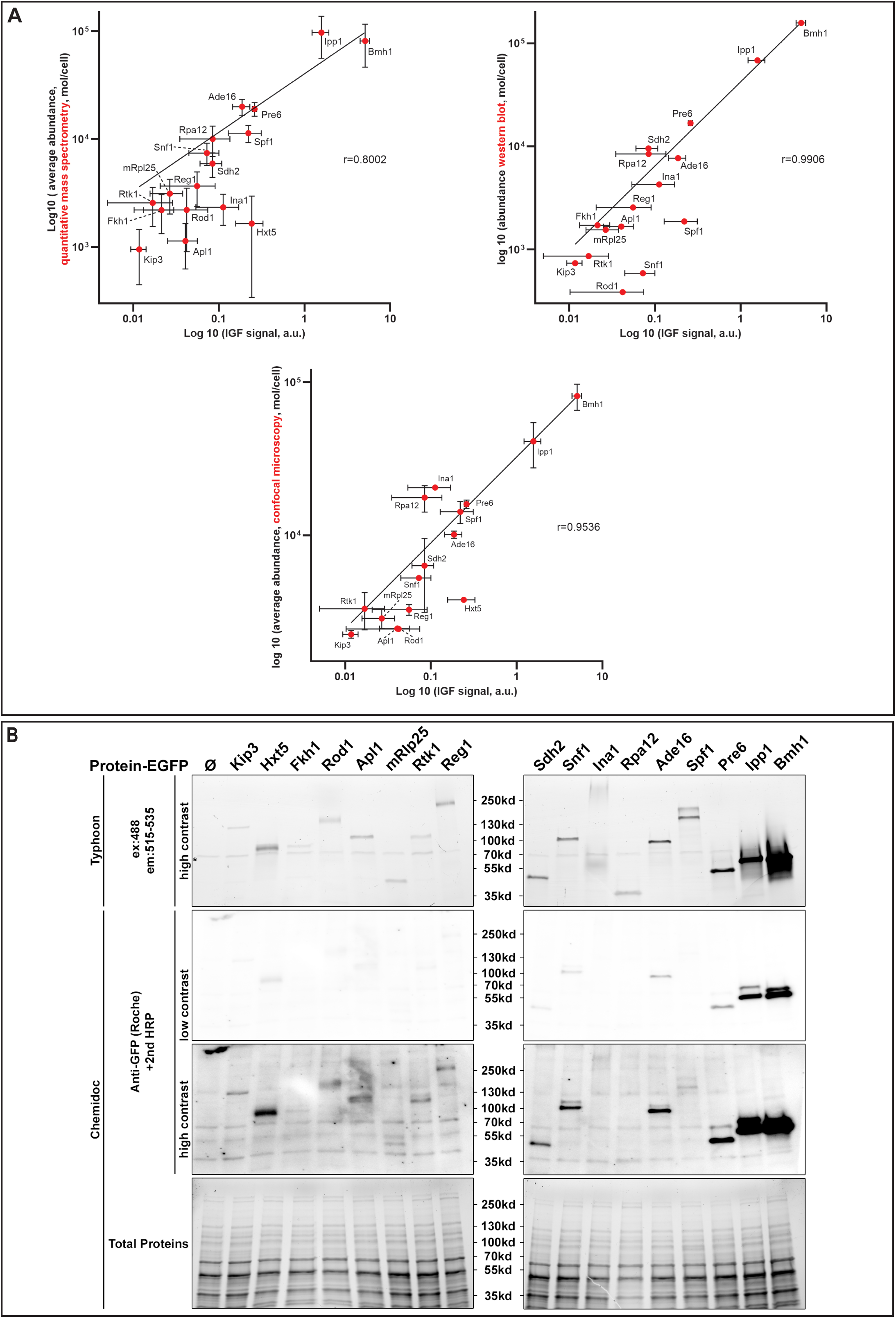
Correlation of IGF signals with published abundances for various proteins and comparison with chemiluminescence detection. A. Correlation of IGF intensities with published abundance based on quantitative mass spectrometry experiments, western blot experiments, of confocal microscopy experiments. Data were from (Ho et al., 2018) and retrieved from SGD (www.yeastgenome.org) for untreated/unchallenged cells. B. Comparison of IGF with antibody-based chemiluminescence detection. The gels displayed in Figure 7B (also shown here on the top) were transferred onto a membrane and immunoblotted with the indicated antibodies for comparison.

**Supplementary Figure 9.**
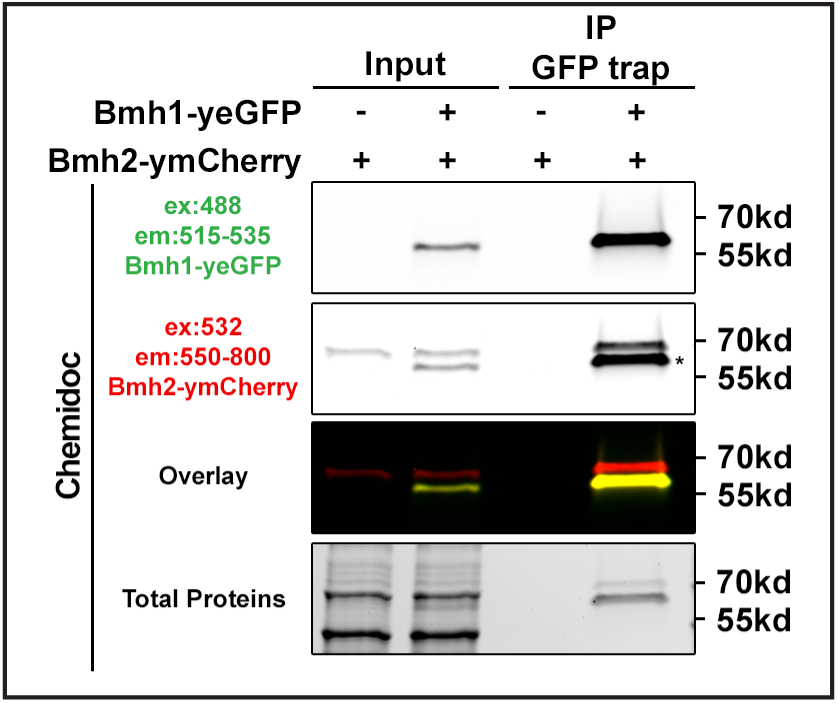
Bleedthrough of green fluorescence in the red fluorescence channel when using a Chemidoc MP. The same gel as that presented in Figure 8A was imaged for red fluorescence in a Chemidoc MP. * indicates the green fluorescent signal that is detected in the red channel.

**Supplementary Figure 10.**
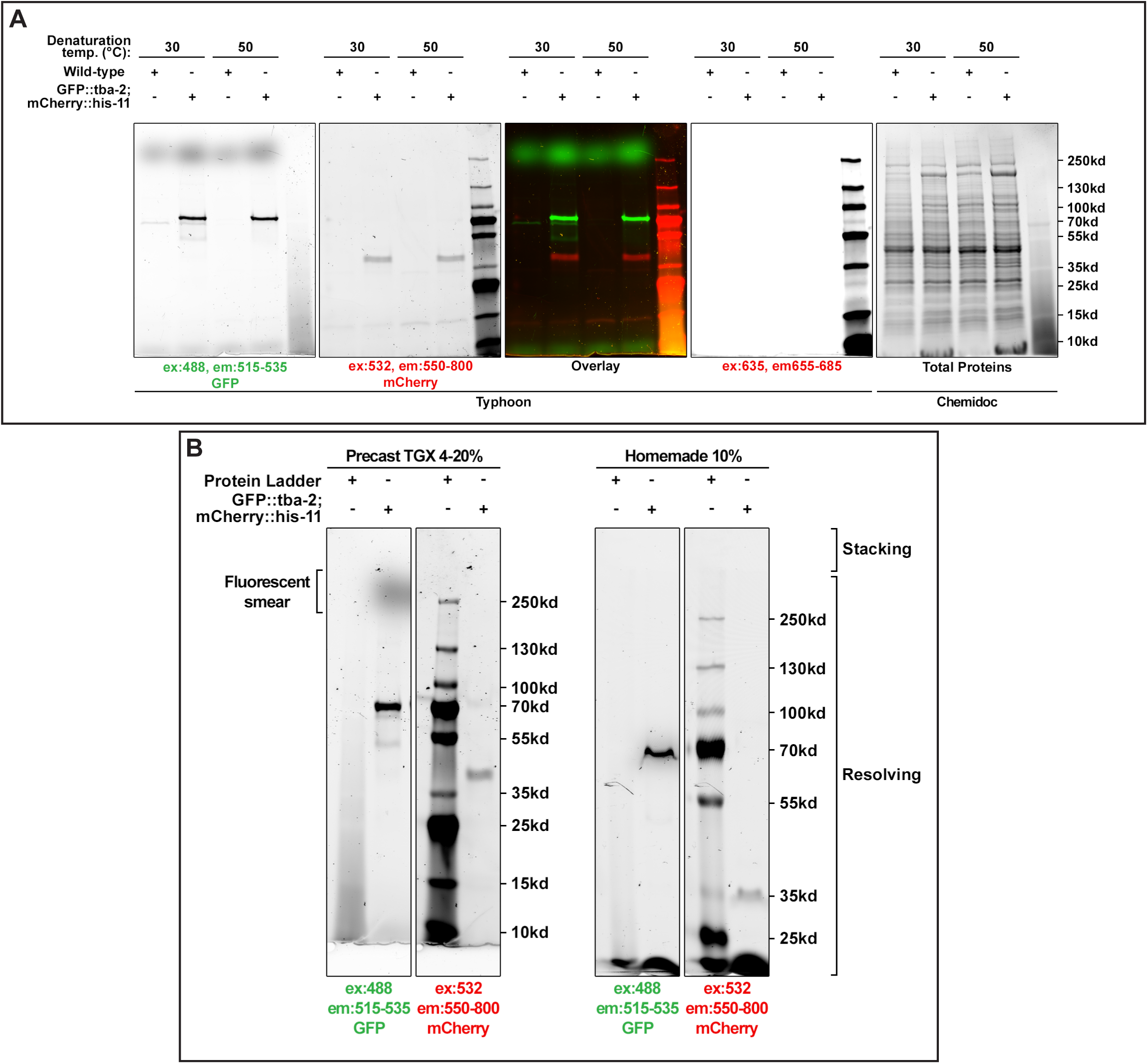
Endogenous fluorescent material in *C. elegans* worm extracts. A. Extended version of the gel presented in Figure 9D. Total protein extracts of *C. elegans* worms were prepared as detailed in *Material and Methods* and loaded onto precast TGX 4-20% gels (Bio-Rad). After migration, gels were imaged for fluorescence using a Typhoon at the indicated excitation/emission wavelengths. A fluorescent smear of unknown origin is visible in the green channel at around 250 kDa. B. The same samples as in A were loaded onto a precast TGX 4-20% gel (Bio-Rad) (left) or homemade 10% SDS-PAGE gel (right). After migration, gels were imaged for fluorescence using a Typhoon at the indicated excitation/emission wavelengths. The fluorescent smear in the green channel at around 250 kDa was no longer observed in homemade gels.

## Notes

### Competing Interest Statement

The authors have declared no competing interest.

### Summary of Updates

The manuscript has been revised following the reviews available from Review Commons. This includes -The addition of an "orange" protein, mKO-kappa, which is more compatible with Chemidoc filters (and as stable as mCherry, making them the two most "resistant" to these conditions) - Figure 5. -Replicated and quantified comparisons between IGF, WB/HRP secondary antibody, and WB/fluorescent secondary antibody. Overall, IGF outperforms in all cases (sensitivity and linearity) - Figure 6. -The study of IGF signal from numerous yeast proteins endogenously tagged with EGFP: different expression levels, various subcellular localizations, cytosolic or membrane proteins, in oligomeric complexes (proteasome, ribosome, RNA Pol) or as monomers. This was conducted to address reviewers' questions: sensitivity, the ability of "mild" denaturing conditions to separate proteins in complexes, behavior of membrane proteins, etc. Data obtained by Matthieu are correlated with published abundances for each protein and compared to a WB anti-GFP. Overall, IGF is applicable to all proteins, including those weakly expressed, membrane proteins, and complexes that are denatured despite the "mild" conditions - Figures 7 and S8. -Compatibility with SNAP labeling (Figure 8C).

